# Human subcutaneous adipose tissue variability is driven by VEGFA, ACTA2, adipocyte density, and ancestral history of the patient

**DOI:** 10.1101/2023.05.31.543052

**Authors:** Megan K DeBari, Elizabeth K Johnston, Jacqueline V Scott, Erica Ilzuka, Wenhuan Sun, Victoria A Webster-Wood, Rosalyn D Abbott

## Abstract

Adipose tissue is a dynamic regulatory organ that has profound effects on the overall health of patients. Unfortunately, inconsistencies in human adipose tissues are extensive and multifactorial including large variability in cellular sizes, lipid content, inflammation, extracellular matrix components, mechanics, and cytokines secreted. Given the high human variability, and since much of what is known about adipose tissue is from animal models, we sought to establish correlations and patterns between biological, mechanical, and epidemiological properties of human adipose tissues. To do this, twenty-six independent variables were cataloged for twenty patients that included patient demographics and factors that drive health, obesity, and fibrosis. A factorial analysis for mixed data (FAMD) was used to analyze patterns in the dataset (with BMI > 25) and a correlation matrix was used to identify interactions between quantitative variables. Vascular endothelial growth factor A (VEGFA) and actin alpha 2, smooth muscle (ACTA2) gene expression were the highest loading in the first two dimensions of the FAMD. The number of adipocytes was also a key driver of patient-related differences, where a decrease in the density of adipocytes was associated with aging. Aging was also correlated with a decrease in overall lipid percentage of subcutaneous tissue (with lipid deposition being favored extracellularly), an increase in transforming growth factor-β1 (TGFβ1), and an increase in M1 macrophage polarization. An important finding was that self-identified race contributed to variance between patients in this study, where Black patients had significantly lower gene expression levels of TGFβ1 and ACTA2. This finding supports the urgent need to account for patient ancestry in biomedical research to develop better therapeutic strategies for all patients. Another important finding was that TGFβ induced factor homeobox 1 (TGIF1), an understudied signaling molecule, is highly correlated with leptin signaling and was correlated with metabolic inflammation. Finally, this study revealed an interesting gene expression pattern where M1 and M2 macrophage markers were correlated with each other, and leptin, in patients with a BMI > 25. This finding supports growing evidence that macrophage polarization in obesity involves a complex, interconnecting network system rather than a full switch in activation patterns from M2 to M1 with increasing body mass. Overall, this study reinforces key findings in animal studies and identifies important areas for future research, where human and animal studies are divergent. Understanding key drivers of human patient variability is required to unravel the complex metabolic health of unique patients.

## 2. Introduction

Long thought of as biologically inert, white adipose tissue is now recognized as a highly active endocrine organ that plays a critical role in systemic hormone, cytokine, immune, and metabolic regulation. Given its dynamic state and regulatory role in the body, changes in adipose tissue have profound effects on the health of patients. For example, expansion of adipose tissue in obesity is a risk factor for a plethora of diseases from diabetes to certain cancers [1–2]. Furthermore, inconsistencies in human adipose tissues are extensive and multifactorial including large variability in cellular sizes, lipid content, inflammation, mechanics, extracellular matrix components, and cytokines secreted [3–6]. This large patient variability leads to poor fat graft consistency and survival when the tissue is implanted [7–10] and inadequate treatment options for disorders related to adipose tissue dysfunction. For example, differences in patient demographics such as race and bariatric surgery have led to profound differences in insulin sensitivity between patients [11–16]. Therefore, determining key variables that drive the large variability in human adipose tissue structure and function is critical.

The predominant cell in adipose tissue is the adipocyte, which stores triglycerides that can be released into the blood stream when energy is required. Additional cells make up the stromal vascular fraction (SVF), which is composed of adipose-derived stem cells (ASCs), preadipocytes, immune cells, fibroblasts, pericytes and endothelial cells [17]. White adipose tissue hosts a variety of resident and infiltrating immune cells, including T and B lymphocytes, macrophages, neutrophils, and eosinophils, that maintain tissue homeostasis [18].

From obese animal models it is known that adipose tissue acts as a reservoir, buffering nutrient surplus with lipid storage and tissue expansion through adipocyte enlargement (hypertrophy) and formation of new adipocytes (hyperplasia) [19–20]. While adipocyte hypertrophy enables adipose tissue to store more lipids, protecting other organs from lipotoxic stress, it decreases the surface area to volume ratio, resulting in ineffective nutrient transport, and poor cell signaling promoting tissue dysfunction and metabolic disorders [21]. In fact, hypertrophic adipocyte growth correlates with diabetes in obese humans [22] whereas hyperplastic growth correlates with improved insulin sensitivity in human subjects [23].

To limit adipose tissue expansion transforming growth factor-β1 (TGFβ1) acts as an inhibitor of adipogenesis [24] and drives fibrotic changes in adipose tissue [25]. Adipose tissue fibrosis is defined as the chronic exposure of adipose tissues to inflammation and hypoxia that leads to a state of extracellular matrix remodeling and collagen deposition [26]. Obesity is one of the main factors leading to fibrosis of adipose tissue [27–28]; however not every patient that is obese will have fibrotic tissue.

Since much of what we know about adipose tissue is from animal models, in this study we sought to establish correlations and patterns between biological, mechanical, and epidemiological properties of human adipose tissues through statistical analyses (**Figure 1**). Based on key roles in health, obesity, and fibrosis 20 independent variables were chosen. The independent variables were paired with six categories of patient demographics (**Table 1**) that were available for each sample. Here, the ultimate goal was to establish key variables that drive patient variability and to determine if similar findings in human tissues are consistent with those of animal studies.

**Figure 1.**
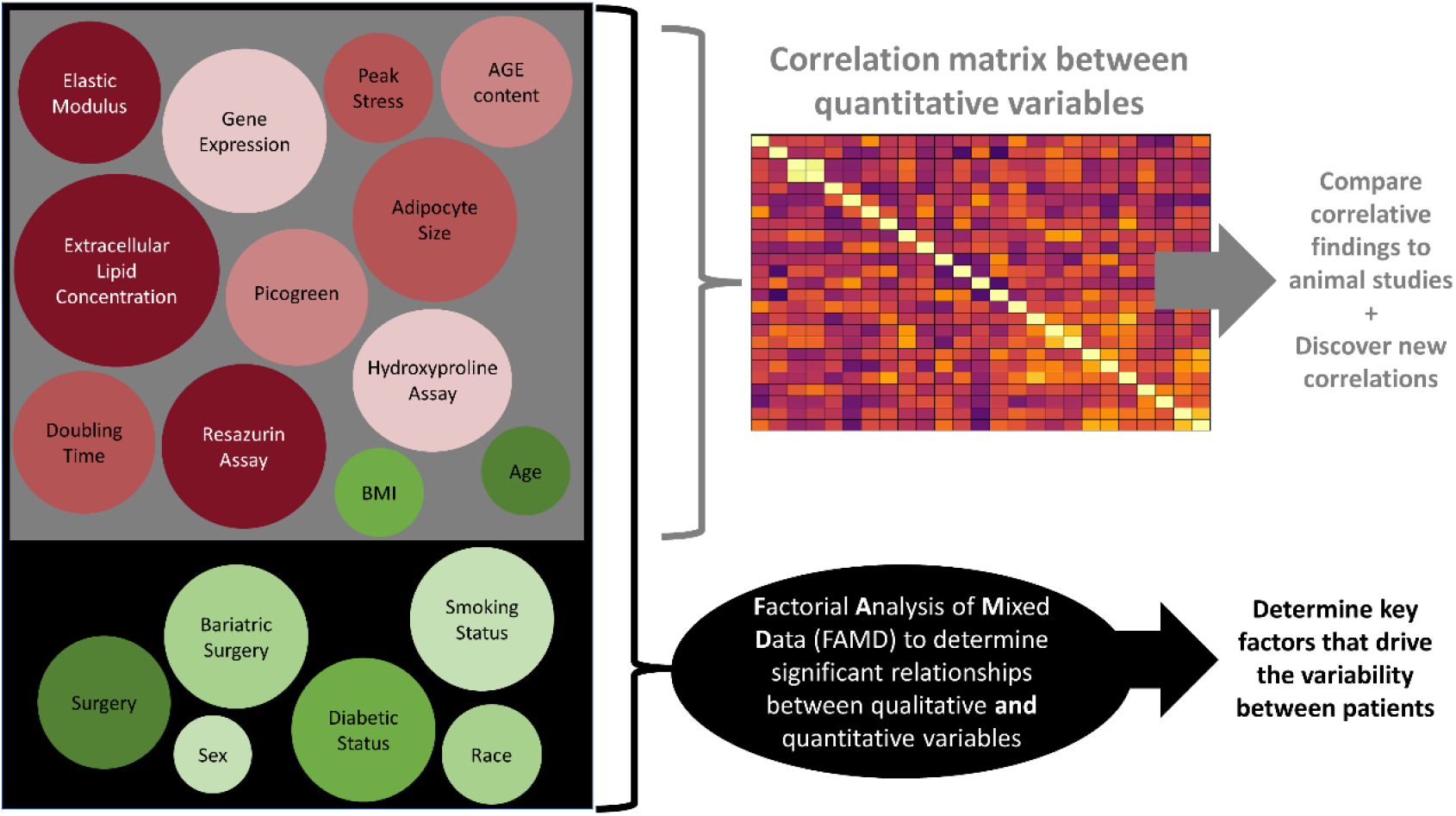
Study design showing how patient demographics (green) and tissue characterization techniques (red) fed into two different statistical models where: 1) each quantitative variable (grey box) was plotted in a correlation matrix against the other quantitative variables to determine correlative relationships and 2) qualitative and quantitative variables (all of the variables in the black box) were fed into a Factorial Analysis of Mixed Data (FAMD) to determine significant relationships between the variables. Correlations were used to compare the human data to published data from animal models (i.e. In animals, TGFβ1 gene expression is related to enhanced collagen deposition, is it the same in humans?) and to discover new correlations (i.e. There is a correlation between gene expression between TGIF and leptin). The FAMD indicated what variables account for the most variation in the dataset (with BMI>25).

**Table 1.** Breakdown of patient demographics used in this study.

| Demographic |  | Number of patients |
| --- | --- | --- |
| **Race** | Black | 6 |
|  | White | 14 |
| **Sex** | Female | 17 |
|  | Male | 3 |
| **Surgery** | Abdominoplasty | 5 |
|  | Panniculectomy | 15 |
| **Bariatric Surgery** | Previous Bariatric Surgery | 13 |
|  | No Bariatric Surgery | 7 |
| **Diabetic Status** | Diabetic | 5 |
|  | Formerly Diabetic | 2 |
|  | Not Diabetic | 13 |
| **Smoking Status** | Non-Smoker | 11 |
|  | Smoker | 9 |

## 3. Methods

### 3.1. Adipose Tissue Procurement

Samples from 20 patients were used in this study. To randomly select patients, adipose tissue was requested only knowing the surgery type. All patient demographics available were used in analysis of this study, including gender, body mass index (BMI), diabetic status, smoking status, age, surgery the adipose tissue was collected from, previous weight loss strategy, and race (**Table 1**). Other information, such as comorbidities was not provided for all patients so this information was excluded from this study but would be advantageous to include in the future. It is important to note, that while the patient’s diabetic status was provided, the type (type I or type II) was not disclosed. Additionally, because the quantity, duration, and quit date was not provided for patients identified as former smokers they were grouped with current smokers. This grouping was done to account for residual or long-lasting effects of smoking. Future work should explore this variable further. Adipose tissue was procured from the Adipose Stem Cell Research Center at the University of Pittsburgh’s Medical Center’s Department of Plastic Surgery. Due to the availability of samples, only overweight and obese patients were obtained (BMI >25). Subcutaneous adipose tissue samples were dissected from abdominal fat taken from either an abdominoplasty or a panniculectomy surgery. A bulk sample with a minimum size of 4” by 4” was used for each experiment. For all testing, samples were taken from directly below the fascia of scarpa.

### 3.2. Imaging

Tissue samples were isolated from bulk tissue in a 3×3 grid. The tissue samples were fixed in formalin (Sigma-Aldrich, St. Louis, MO). 6 of the samples were randomly selected and stained with AdipoRed (Lonza, Walkersville, MD) (1:35) and Alexa Fluor Phalloidin 488 (Thermo Fisher Scientific, Waltham, MA) (1:100) and imaged using confocal and multiphoton microscopy (Nikon, Melville, NY) to visualize lipids, f-actin, and collagen *via* Second Harmonic Generation. Images taken using confocal microscopy were used to determine the size of extracellular lipid droplets. 2 samples per patient were imaged, with 2 images being captured per sample for a total of 80 images. A total of 1904 extracellular lipid droplets were measured. Extracellular lipid droplets were defined as lipids that were not intracellularly contained, as indicated by the exclusion from the cytoskeletal Phalloidin staining. The mean diameter was determined to be 7.27 μm, with a standard deviation of 8.16 μm (**Figure 2**). To encompass a majority of extracellular lipid droplets the mean plus one standard deviation (15.43 μm) was determined to be the maximum size of what we defined as an extracellular lipid droplet. This accounted for 84.1% of extracellular lipid droplets in the samples imaged and was well-below the smallest adipocytes which are >20 μm [3]. Every patient and field of view had extracellular lipid droplets, however the number of lipid droplets varied slightly from patient to patient.

**Figure 2.**
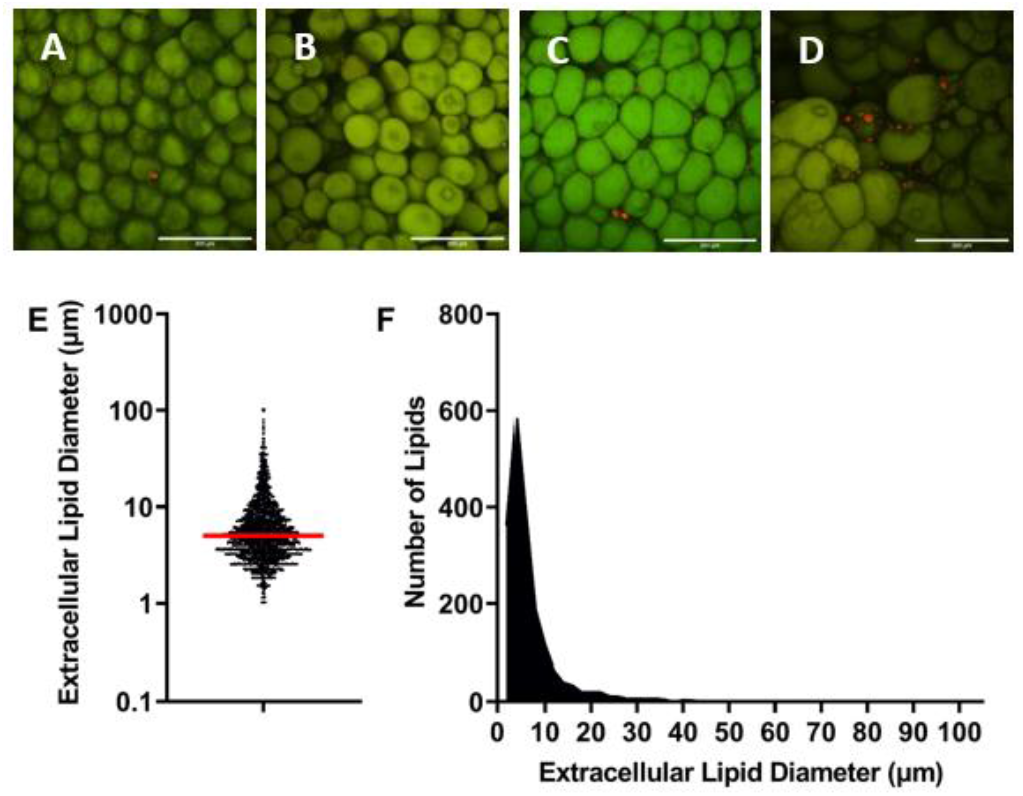
(A-D) Representative images used to analyze extracellular lipid diameter. Samples were stained with Adipored (red) and Phalloidin 488 (green). Extracellular lipids appear red while adipocytes (stained by both Adipored and Phalloidin 488) appear green. Scale bars are 200 μm. Each image is from a different patient. (E) Each dot on the graph represents an extracellular lipid droplet diameter measurement, with the mean represented by the red line (mean – 7.27 μm, standard deviation – 8.16 μm). (F) Histogram showing frequency of extracellular lipids with specific diameters, where the majority of extracellular lipids measured were under 15 μm. n=1904.

The extracellular and intracellular lipid droplets and collagen were visualized with multiphoton microscopy. Depending on the sample and the location of the image the collagen to adipocyte distribution varied. To account for the high variability of collagen/adipocyte distribution many images were captured and analyzed. 6 samples were imaged, with 3 images being captured per sample for a total of 360 images. These images were used to evaluate: 1) adipocyte diameter; 2) the extracellular lipid percent; and 3) whether extracellular lipids were in high-density collagen areas (fibrotic regions). Only lipid droplets that could be clearly measured were used for analysis. The number of adipocytes and extracellular lipid droplets were counted and used to determine the extracellular lipid percent defined as:

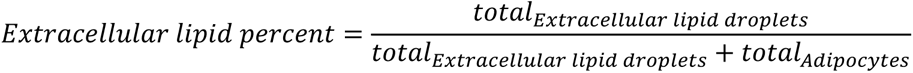

Where total _Extracellular lipid droplets_ represents the total number of extracellular lipid droplets in each image and total _Adipocytes_ represents the total number of adipocytes counted in each image. Additionally, the location of the extracellular lipid droplets was categorized by their proximity to collagen. Extracellular lipid droplets embedded in, or touching collagen fibrils were identified as “extracellular lipid droplets in collagen.” This was used to determine if extracellular lipid droplets were co-localized within the collagen:

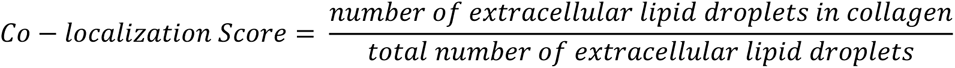

All image analysis measurements were done using ImageJ. An average of 528 adipocytes were measured per patient to obtain diameter metrics. All images were taken using the same magnification. For full set of quantification results for each patient see **Supplement 1**.

### 3.3. DNA Content

Picogreen assays (Thermo Fisher Scientific, Waltham, MA) were performed on each patient’s tissue samples following the manufacturer’s procedure to assess DNA content. The DNA content was normalized to the weight of each tissue sample. Samples were run in technical duplicates. N=5 for each patient.

### 3.4. Metabolic Activity / Redox readout

Resazurin (Thermo Fisher Scientific, Waltham, MA) was diluted to 1 mM with phosphate buffered saline (PBS)(pH 7.4). This was diluted further to 0.05 mM solution using cell culture media (DMEM with 10% FBS and 1% Pen-Strep). 1 mL of the 0.05 mM resazurin/media solution was placed in a 24-well plate with the tissue samples. The samples were incubated at 37°C for 2.5 hours. Using a microplate reader (SpectraMax i3x), the absorbance at 570/600 nm was measured. Absorbance data were normalized to the average DNA content per gram of tissue for each patient. Samples were run in technical duplicates. N=5 for each patient.

### 3.5. Collagen Content

Hydroxyproline assays (Sigma-Aldrich, St. Louis, MO) were performed on each patient’s tissue samples following the manufacturer’s protocol to assess collagen content. Briefly, approximately 10 mg was homogenized in 100 µL of water (exact tissue weight was recorded before homogenization). 100 µL of 12 M HCL was added and hydrolyzed for 3 hours. The supernatant was then transferred to a well place and allowed to evaporate. Chloramine T/Oxidation buffer and Diluted DMAB Reagent were then added following the manufacturer’s protocol. Collagen content was normalized to the weight of the initial tissue sample. Samples were run in technical duplicates. N=1 for each patient.

### 3.6. Advanced Glycation End Products (AGE)

AGE assays (Abcam, Cambridge, MA) were performed on each patient’s tissue samples. Tissue samples were flash frozen in liquid nitrogen and stored at −80°C until use. Tissue samples were then suspended in PBS (pH 7.4) (100 mg of tissue to 1 mL of PBS), homogenized, and stored at −20°C overnight. The following day, homogenized tissues were subjected to two freeze thaw cycles. 1 mL of the solution was then centrifuged at 5000 x g at room temperature for 5 min. Samples for this assay were taken from right below the lipid layer of the centrifuged homogenized tissue to avoid the lipid layer. The assay was performed following the manufacturer’s procedure. Samples were run in technical duplicates. N=1 for each patient.

### 3.7. Compression Testing

Tissue samples were compressed to 90% strain at a rate of 1 mm/min using an electromechanical universal testing system (MTS Criterion, MTS, Eden Prairie, MN). The elastic modulus was determined by the linear portion of the stress strain curve between 60-90% strain. The adipose tissue samples had stress-strain curves similar to elastomers [29]. The stress remained low until around 50-60% strain when it began to increase drastically. 60-90% strain was used to determine the moduli because this is the region of the curve where the extracellular matrix proteins (ECM) are being compressed. Prior to the drastic increase in stress, it is thought that adipocytes are being compressed, but not to failure. Once the adipocytes are compressed to failure, the stress increases quickly. This is further supported visually by the lipids or “oil” being seen pooling around the sample as the stress began to increase. Each sample was compressed to 90% strain and at this point the peak stress was recorded. N= 10 for each patient.

### 3.8. Stromal Vascular Fraction (SVF) Doubling Time

The SVF was isolated from subcutaneous adipose tissue as described previously [30]. The cells were isolated by mechanically blending the adipose tissue until the texture resembled lipoaspirate. The tissue was then incubated in a collagenase solution (0.1% collagenase, 1% bovine serum albumin, 98.9% phosphate buffer solution) (Thermo Fisher Scientific, Waltham, MA) at a 1:1 ratio for 1 hour at 37°C. Following the incubation, the solution was centrifuged (5 minutes at 300 g) to isolate the SVF. The cells were resuspended in media (DMEM with 10% FBS and 1% pen-strep), centrifuged, and seeded into flasks. Once cells reached confluency, cells were trypsinized and 10,000 cells/well were seeded into a 6-well plate to ensure comparable log-phase growth. Media was changed every 2-3 days. 3 images of each well were taken every other day until confluency was reached. Cells were counted and the doubling time was calculated using the following equation:

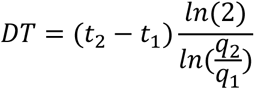

Where DT = doubling time, t_2_ = second time point, t_1_ = first time point, q_2_ = number of cells at second time point, and q_1_ = number of cells at first time point. N=5 for each patient. Due to a COVID19 laboratory shutdown, patients 5 and 6 are missing doubling time data. For the complete set of data for each patient related to mechanical properties, collagen content, AGEs, metabolic activity, and doubling rate see **Supplement 2**.

### 3.9. Reverse Transcription Quantitative Real-Time Polymerase Chain Reaction (RT-qPCR)

Tissue samples were flash frozen in liquid nitrogen and stored at −80°C until RNA isolation. Total RNA was isolated using the RNeasy Lipid Tissue Mini Kit (QIAGEN, Venlo, Netherlands) and concentration was measured using the NanoDrop 2000c spectrophotometer (Thermo Scientific, Waltham, MA). RNA was processed into cDNA using the iScript cDNA Synthesis Kit (Bio-Rad, Hercules, CA). qPCR reactions were performed with 100 ng of cDNA per reaction. Succinate dehydrogenase complex flavoprotein subunit A (SDHA) (ID: 6389) was used as a reference housekeeping gene [31] and interleukin 6 (IL6) (ID: 3569), tumor necrosis factor α (TNFα) (ID: 7124), TGFβ induced factor homeobox 1 (TGIF1) (ID: 7050), leptin (LEP) (ID: 3952), adiponectin (ADIPOQ) (ID: 9370), transforming growth factor β 1 (TGFβ1) (ID: 7040), actin alpha 2, smooth muscle (ACTA2) (ID: 59), CD163 (ID: 9332), CD86 (ID: 942), and vascular endothelial growth factor A (VEGFA) (ID: 7422) were included as genes of interest. All patient and primer combinations were performed with technical duplicates. The CT value was determined through regression analysis performed by the Bio-Rad CFX96 Real-Time System and C1000 Touch Thermal Cycler (Bio-Rad, Hercules, CA). Since there was no control group only ΔCT was calculated:

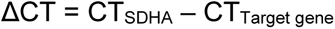

Using this formula higher CT values of the target gene, which indicate lower gene expression, would have lower ΔCT values. A CT of 45 was assigned for genes that were not detected after 40 cycles. For the full set of patient PCR results see **Supplement 3**.

### 3.10. Statistics

Quantitative variables were evaluated for differences between patients using either a one-way ANOVA or two-way ANOVA followed by a Tukey’s post-hoc analysis (GraphPad Prism 9.0.0). A one-way ANOVA was used to determine statistical differences between adipocyte diameters, number of extracellular lipid droplets in collagen/total number of extracellular lipid droplets co-localization, elastic moduli, peak stress, doubling time, and metabolic activity. A two-way ANOVA was used to determine statistical differences between the percent of pixels (collagen/lipids). Comparisons between ancestry of the cells (White versus Black) were performed with a t-test. P values less than 0.05 were determined statistically significant. All data is represented as mean +/− standard deviation. A factorial analysis for mixed data (FAMD) was performed on the data using R statistics software. To get all of the variables on the same scale, data was standardized before conducting the analysis by subtracting the mean of the data set from every value and then dividing by the standard deviation [32–33]. Because complete data sets are required to perform a FAMD, the missMDA package was used to estimate the two missing doubling time data points [34] from a COVID19 lab closure. The FAMD analysis was performed using the FactoMineR package, with additional graphs being created with the factoextra package. Cos2 values were normalized to the highest value in each dimension to determine significant correlations. Coordinates for quantitative and qualitative data for each dimension are included in **Supplement 4**. R-code used to generate FAMD results is included in **Supplement 5**. Using the GraphPad Prism statistical software, all quantitative data was used to generate a correlation matrix using the Pearson correlation coefficient with a confidence interval of 95%.

## 4. Results & Discussion

Dimensionality reduction is used to identify relationships amongst variables and extract patterns in datasets. To determine if there were relationships between variables in our dataset, we used two different statistical models: 1) a correlation matrix to evaluate relationships between quantitative variables (**Figure 3**) and 2) a factorial analysis of mixed data (FAMD) to understand the relationships between qualitative (race, sex, surgery type, diabetic status, smoking status) and quantitative variables. A FAMD is similar to principal component analysis (quantitative data only) and multiple correspondence analysis (qualitative data only), but both quantitative and qualitative data can be analyzed [34]. With the FAMD, the number of dimensions needed to account for > 80% of the variability in our dataset was determined to be 9 (**Table 2**, for full set of coordinates see **Supplement 4**). The largest loading was the variable that contributed the most variance in the given dimension, meaning that variability in these key factors were correlated with variability in the rest of the dataset and are essential contributors to patient-related differences.

**Figure 3.**
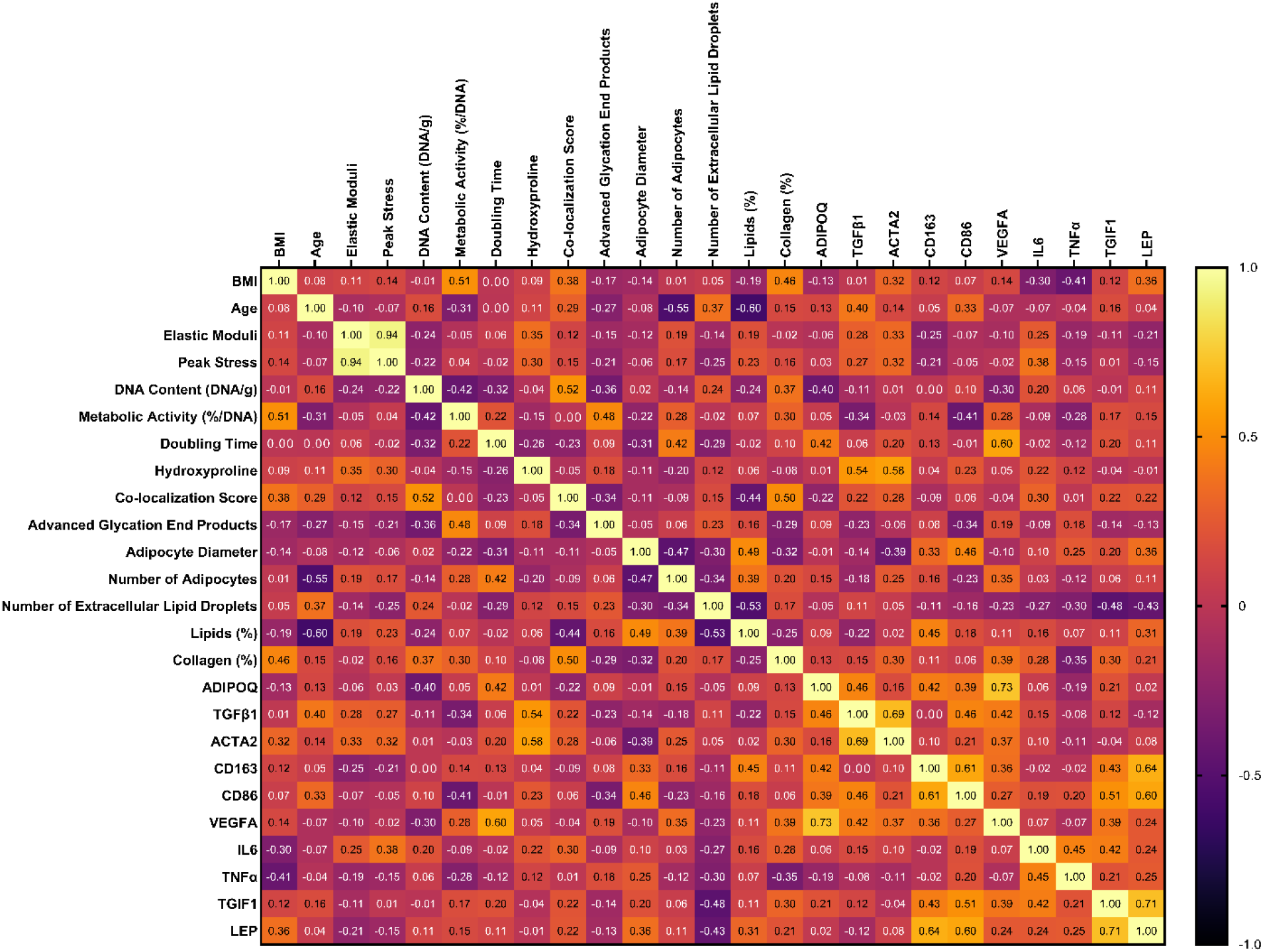
Amongst quantitative variables, many correlations exist in BMI, age, mechanical properties, cell number (DNA content), metabolic activity, doubling time of the stromal vascular fraction, collagen content (hydroxyproline), advanced glycation endproducts, morphological measurements, histological quantification, and gene expression. The correlation matrix indicates the correlation between quantitative variables (samples from N=20 patients with BMI>25). A value greater than 0.7 is considered a high correlation, 0.5-0.7 is considered a moderate correlation, and 0.3-0.5 is considered a low correlation.

**Table 2.** Eigenvalues and variance determined through factorial analysis of mixed data (FAMD) analysis.

| *Dimension* | *Eigenvalue* | *Variance (%)* | *Cumulative Variance (%)* | *Largest Loading* |
| --- | --- | --- | --- | --- |
| 1 | 4.89 | 15.29 | 15.29 | VEGFA |
| 2 | 4.28 | 13.38 | 28.67 | ACTA2 |
| 3 | 3.68 | 11.5 | 40.17 | Number of Adipocytes |
| 4 | 3.52 | 11.01 | 51.18 | Race |
| 5 | 3.05 | 9.52 | 60.7 | Elastic Moduli |
| 6 | 2.26 | 7.08 | 67.78 | Diabetic Status |
| 7 | 1.89 | 5.91 | 73.69 | Diabetic Status |
| 8 | 1.84 | 5.75 | 79.44 | Smoking Status |
| 9 | 1.53 | 4.78 | 84.22 | Advanced Glycation End Products |

### 4.1 Levels of Vascular Endothelial Growth Factor A (VEGFA) gene expression accounted for the most variance in the full dataset of variables (qualitative and quantitative)

The variable that contributed the most to the first dimension of the FAMD analysis was gene expression of VEGFA (**Table 2**). This means that variability in VEGFA gene expression was linked to the most patient-related differences in the other variables in this study. In all individuals, the resident vasculature of adipose tissue acts to ensure adequate blood flow and nutrient/waste transport to allow for adipose tissue expansion and metabolism. In adipose tissue, VEGFA is a pleiotropic molecule that is involved in vasculogenesis, angiogenesis, vascular permeability, tissue remodeling, and metabolic effects (see our recent reviews: [35–36]). Mouse models that overexpress VEGFA result in an increase in adipose tissue vascularization, systemic protection against metabolic dysfunction from a high-fat diet, and higher energy expenditure [37–38]. In the patient population evaluated here, higher gene expression of VEGFA was positively correlated with a longer doubling time of seeded stromal vascular fraction cells (0.60) and higher adiponectin gene expression (0.73) (**Figure 4**).

**Figure 4.**
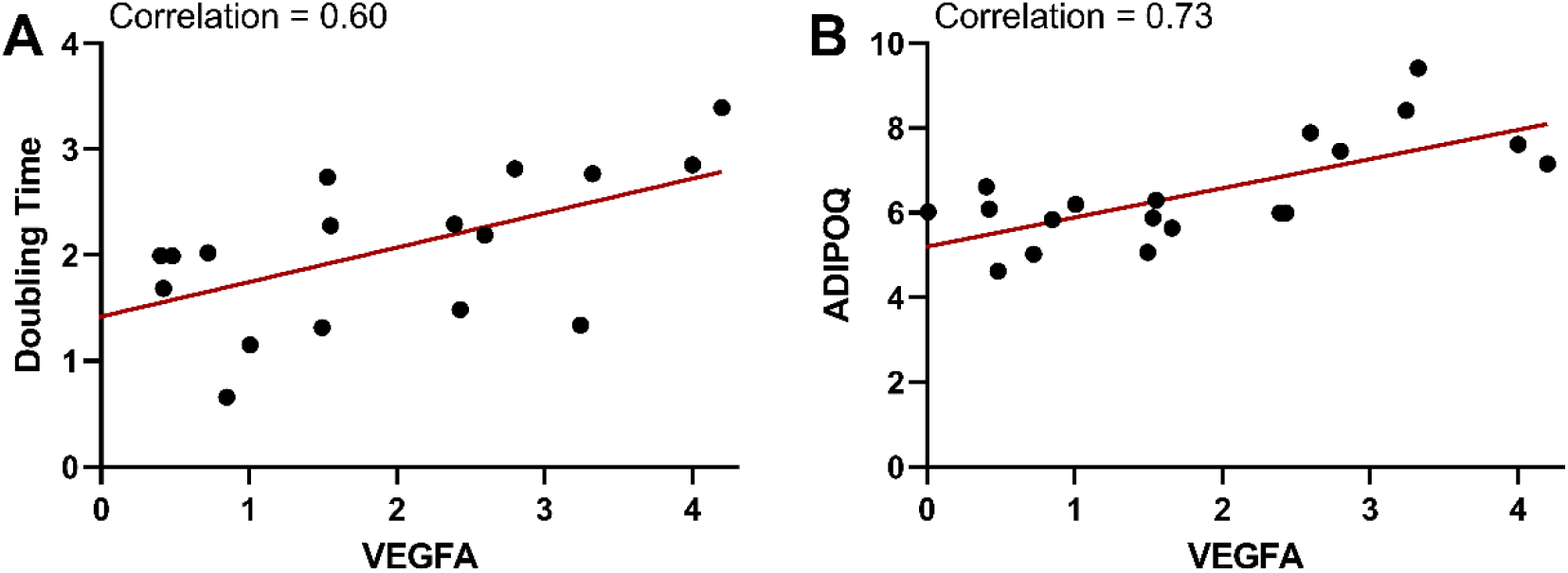
Gene expression of VEGFA was correlated to stromal vascular doubling time and adiponectin gene expression. VEGFA gene expression is plotted versus stromal vascular doubling time (A) and adiponectin (ADIPOQ) gene expression (B). Gene expression is represented as a delta CT from the housekeeping (HK) gene (ΔCT = CT_SDHA_ – CT_Target gene_). Therefore, the HK gene is equal to 0 on the plots. With the formula used, gene expression is relative to the housekeeping gene, and increases at higher values. Correlations from the correlation matrix are indicated on each plot.

VEGFA is widely expressed in adipose tissue by both the SVF as well as the mature adipocytes [39]. When the SVF is expanded *in vitro* the predominant cells that persist are multilineage ASCs [40]. Therefore, the doubling time in our study reflects the proliferation of the ASCs. Many interacting factors can impact the proliferative capacity of ASCs, for example, diabetes [41], aging [42–43], and certain drugs [44]. However, the correlation between VEGFA and longer ASC doubling time has not been reported. It is well accepted that while proangiogenic, the exogenous application of VEGFA on endothelial cells can have a preferential response to either proliferation or migration depending on the need [45]. One possibility is that similar to endothelial cells, ASCs may have a more migratory response to support angiogenesis. Additionally, in adipose tissue the balance between stem cell self-renewal and differentiation must be maintained. The regulatory mechanisms for maintaining the stem cell state as well as fate determination are highly conserved. With how heavily intertwined angiogenesis and adipogenesis are it is not surprising that VEGF is one of these regulators as blocking VEGFR2, preadipocyte differentiation is also inhibited [46]. Since VEGFA is increased during adipocyte differentiation [47], this positive correlation could indicate that the individual is experiencing more hyperplastic adipose tissue growth where ASCs are recruited and differentiate towards adipocytes rather than undergoing proliferation. Supporting this theory, a longer doubling time was also correlated with a higher number of adipocytes (**Supplemental Figure 2B,** 0.42).

The strong correlation between adiponectin and VEGFA supports findings that adiponectin is down-regulated in VEGFA ablated mice [48–49]. Adiponectin is often considered to be a proangiogenic factor [50–51]. In certain diseases such as chondrosarcoma [52] and rheumatoid arthritis [53], adiponectin promotes VEGFA-dependent angiogenesis. However, other studies have suggested adiponectin induces endothelial cell apoptosis, reduces their proliferative capacity [54], and inhibits endothelial migration [55]. Therefore, further research is required to discern the correlation observed between adiponectin and VEGFA in the patient population evaluated here. VEGFA also is known to have a protective role in adipose tissue inflammation by recruiting M2 macrophages to adipose depots in mice [56]. Supporting this finding, our dataset indicated there was a low correlation between VEGFA and CD163 (0.36), with the FAMD indicating these factors were clustered (**Supplement 4**). Overall, these findings support a role of adiponectin and M2 macrophage recruitment in VEGFA-mediated metabolic protection.

### 4.2 TGFβ induced factor homeobox 1 (TGIF1) gene expression was a key contributor to the highest loading direction

The second largest contributor in the first dimension of the FAMD was gene expression values for TGIF1 (**Figure 5**). The role of TGIF1 in adipose tissue is only recently being uncovered. TGIF1 represses TGFβ signaling via multiple pathways including 1) direct Smad2 inhibition forming a transcriptional repressor complex, 2) prevention of Smad2 phosphorylation, or 3) targeting Smad2 for ubiquitin-dependent degradation [57–59]. In preadipocytes, insulin antagonizes TGFβ signaling in preadipocytes through TGIF1 transcription and is required for the differentiation of preadipocytes [60]. In fact, a recent screening study identified lower TGIF1 transcription as a key reducer of lipid accumulation in differentiating stem cells [61]. TGIF1 positively interacted with leptin, IL6, CD163, CD86, and no bariatric surgery, while negatively interacting with the number of extracellular lipid droplets (**Figure 6**).

**Figure 5.**
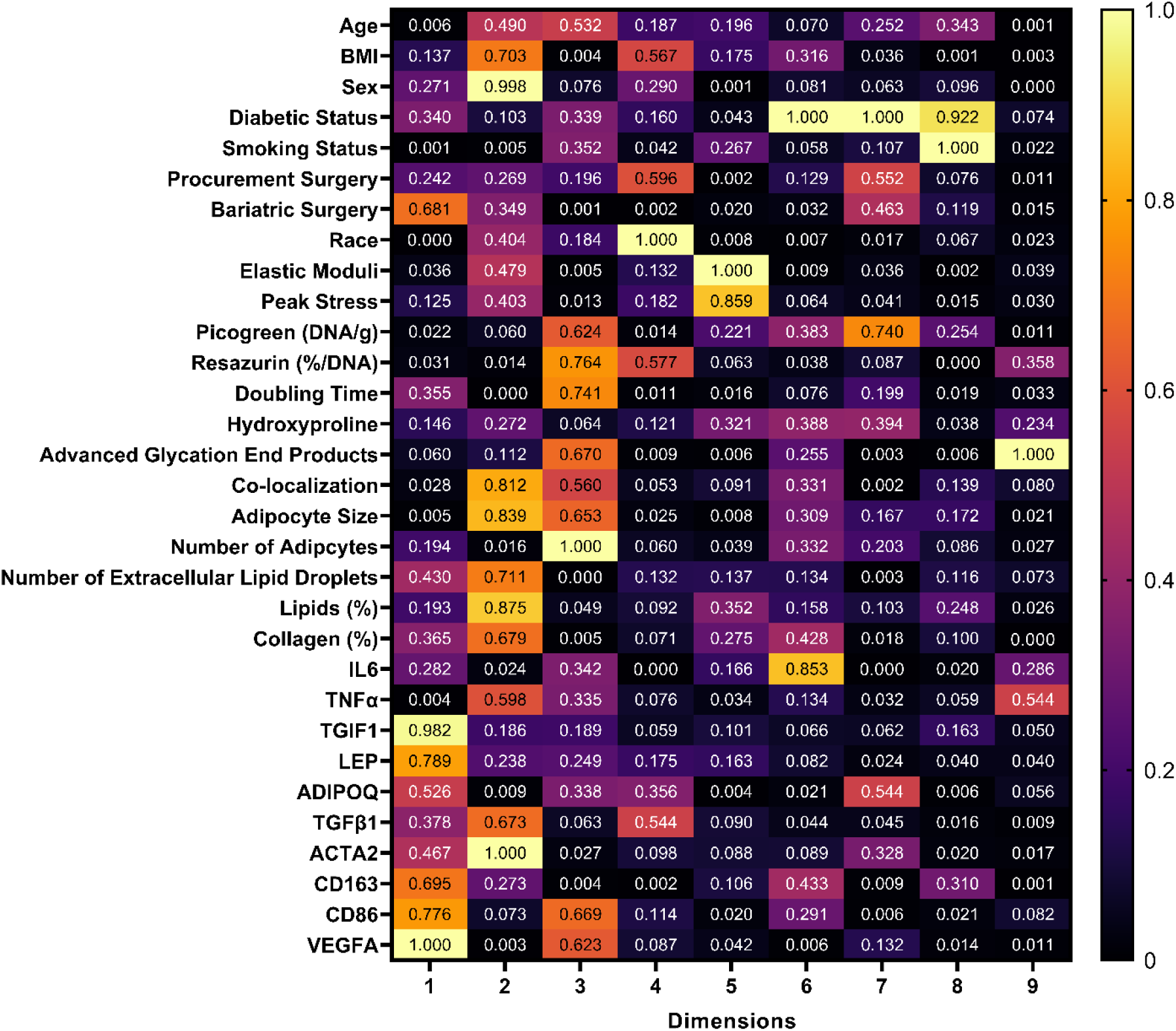
Contribution results generated from the FAMD analysis normalized to the highest contribution in each dimension. In each column the contribution of variables to the 9 dimensions is shown in a heat map, where a value of 1 indicates the largest loading of a variable and the highest contributor in that dimension. For example, in dimension 1 VEGFA is the highest contributor (1.00) followed closely by TGIF1 (0.982).

**Figure 6.**
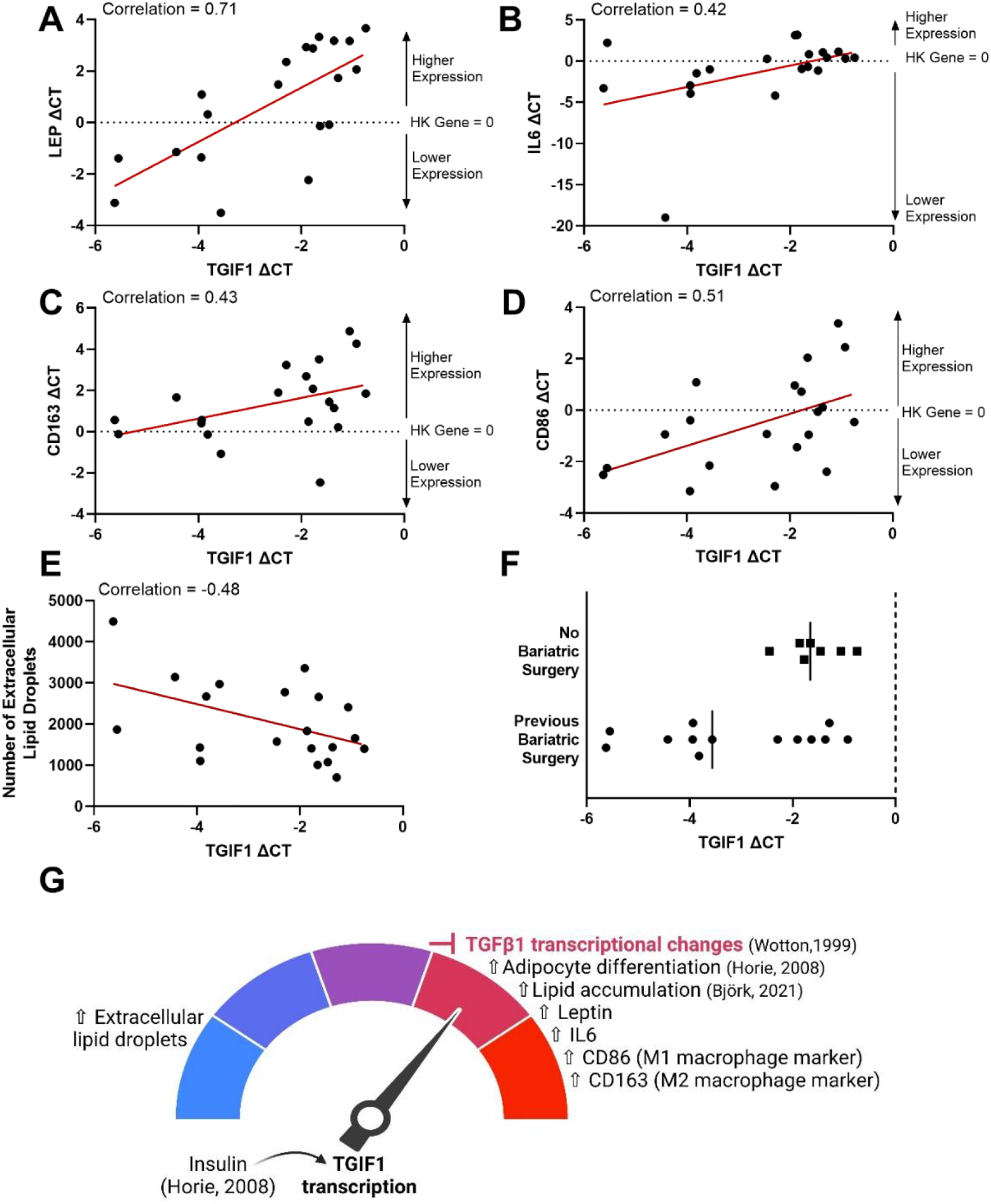
Increased gene expression of transforming growth factor-β induced factor homeobox 1 (TGIF1) was correlated with an increase in gene expression of leptin, interleukin-6. (**IL6), CD163, CD86, and no bariatric surgery, and a decrease in the number of extracellular lipid droplets.** TGIF1 gene expression was plotted versus leptin (LEP) gene expression (A), IL6 gene expression (B), CD163 gene expression (C), CD86 gene expression (D), the number of extracellular lipid droplets counted in histological images (E), and separated by whether the patient had undergone bariatric surgery previously or not. Gene expression is represented as a delta CT from the housekeeping (HK) gene (ΔCT = CT_SDHA_ – CT_Target gene_). Therefore, the HK gene is equal to 0 on the plots. With the formula used, gene expression is relative to the housekeeping gene, and increases from a negative value to a higher positive value. Correlations from the correlation matrix are indicated on each plot. A summary of the results and current literature findings are illustrated (G), where the literature indicates insulin upregulates TGIF1 (Horie, 2008), blocking TGFβ1 transcriptional changes (Wotton, 1999), inducing adipocyte differentiation (Horie, 2008), and increasing lipid accumulation (Bjork, 2021).

Leptin is a pleiotropic hormone predominantly synthesized by adipose tissue that is produced in greater quantities with increasing adipose tissue mass [62]. While leptin is known to stimulate liver fibrosis [63–65], it has been shown to inhibit fibrosis in mouse adipose tissue [66–67]. Our data suggests that TGIF1 could have a role in leptin suppression of fibrosis, as there was a high correlation between higher gene expression levels of TGIF1 and leptin (0.71). It is possible that an increase in leptin stimulates higher TGIF1, suppressing TGFβ-triggered fibrotic changes. However, given the endpoint nature of the study causation cannot be determined. Therefore, determining the directionality of the leptin and TGIF1 interaction will be an important next step. Another striking correlation is that lower levels of TGIF1 are correlated with higher numbers of extracellular lipid droplets (−0.48). Further work needs to explore the role of extracellular lipid droplets observed in adipose tissue and their association with TGIF1.

Higher levels of TGIF1 were positively correlated with Interleukin 6 (Il6) and macrophage surface markers CD163 and CD86 (**Figure 6 B, C and D**, respectively), suggesting high levels of TGIF1 expression could have a role in metabolic inflammation in adipose tissue. Adipocyte secretion of IL6 increases macrophage infiltration in adipose tissue [68] and is linked with reduced insulin sensitivity and diabetes [69]. While both cell surface markers are present on polarized macrophages, CD86 is typically considered to be a classically activated proinflammatory M1 macrophage marker and CD163 is an alternatively activated anti-inflammatory M2 macrophage marker [70–71]. In the overweight/obese population studied here, CD86 and CD163 were positively correlated (0.61). It’s not unusual to see high levels of both CD86 and CD163 gene expression in adipose tissue samples with BMI>25, as others have noted this signature as indicative of metabolic inflammation [71]. Furthermore, TGIF1 has been identified as a potential transcriptional regulator in macrophage activation [72].

The correlation between TGIF1 gene expression and bariatric surgery is interesting. Bariatric surgery results in rapid weight loss and is an effective treatment to restore insulin sensitivity and reduce type 2 diabetes complications [73–74]. Our results indicated that in some patients, TGIF1 levels of patients that had previously undergone bariatric surgery were well-below levels of those that had not (**Figure 6F**). This is an important finding suggesting that some patients that undergo bariatric surgery will have low levels of TGIF1 and will likely not have TGIF1-mediated TGFβ1 signaling suppression. However, lower levels of TGIF1 were not seen in all patients that underwent surgery, suggesting other variables also interact.

It is not surprising that TGIF1 and TGFβ1 gene expression levels were not correlated in our adipose tissue samples (0.12). This is because TGIF1 acts downstream from TGFβ1, repressing the transcriptional response [57] rather than affecting the expression of the TGFβ1 gene. This is seen in other tissues, as well. For example, in the glomerulus, the profibrotic action of TGFβ1 is antagonized by TGIF and blocks α-smooth muscle activation; however high levels of TGIF do not affect TGFβ1-mediated Smad2 phosphorylation and its nuclear translocation [75].

### 4.3 ACTA2 was the second loading dimension of the FAMD

ACTA2 contributed the most to the second dimension of the FAMD. ACTA2 is a gene that encodes smooth muscle alpha (α)-2 actin production which is required for cell movement and muscle contraction. ACTA2 (**Figure 7**) was correlated with TGFβ1 (0.69) and collagen content (0.58), while being inversely related to adipocyte diameter (−0.39). This is consistent with studies where murine adipocytes exposed to a high fat diet upregulate expression of extracellular matrix genes and ACTA2 gene expression characteristic of a myofibroblast-like cell type through reduced PPARγ activity and elevated TGFβ-SMAD signaling [76–77]. The ACTA2 “cellular identity crisis” is thought to drive functionality changes in obese adipose tissues [76] where it is known that enhanced collagen deposition restricts adipocyte size [78]. This was further supported by our data set’s low inverse correlation between collagen percent and adipocyte diameter (−0.32). Collectively, our data suggests a strong role for ACTA2 signaling as a contributor of the variability in human adipose tissues, where high ACTA2 gene expression is related to fibrosis (collagen deposition and TGFβ1) and a reduction in adipocyte cellular diameter.

**Figure 7.**
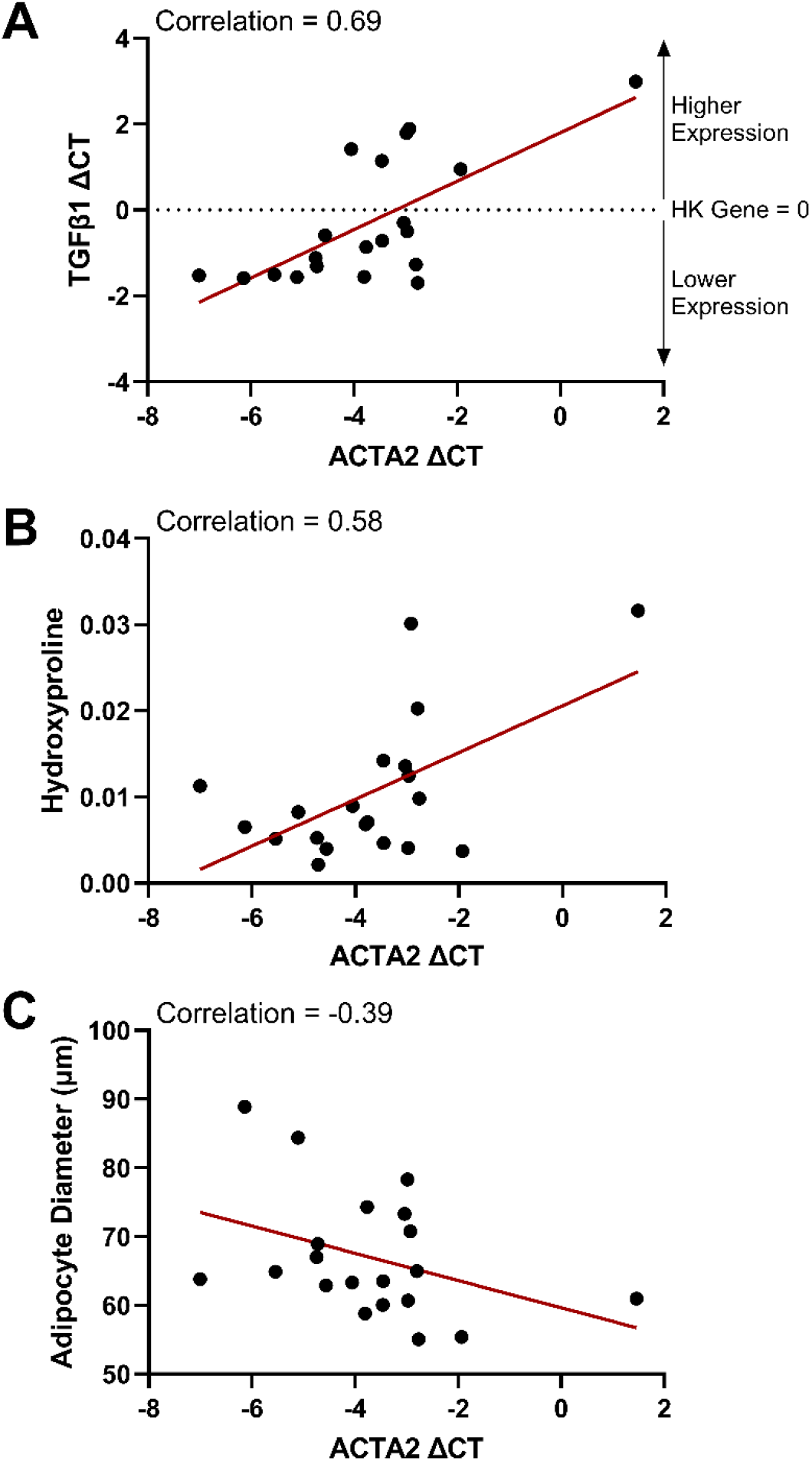
Increased gene expression of actin alpha 2, smooth muscle (ACTA2) gene expression was correlated with an increase in transforming growth factor-β (TGFβ1) gene expression and collagen content (hydroxyproline), (VEGFA), with a corresponding decrease in adipocyte diameter. ACTA2 gene expression was plotted versus TGFβ1 (A), hydroxyproline content (B), VEGFA (C), and measurements of adipocyte diameter in histological images (D), the number of extracellular lipid droplets counted in histological images (E). Gene expression is represented as a delta CT from the housekeeping (HK) gene (ΔCT = CT_SDHA_ – CT_Target gene_). Therefore, the HK gene is equal to 0 on the plots. With the formula used, gene expression is relative to the housekeeping gene, and increases from a negative value to a higher positive value. Correlations from the correlation matrix are indicated on each plot.

### 4.4 TGFβ1 expression

Besides being correlated with ACTA2 (0.69), as discussed in the last section, TGFβ1 was correlated with hydroxyproline content (0.54), the classically activated M1 macrophage surface marker gene expression (CD86, 0.46), and VEGFA gene expression (0.42). These results are consistent with the known fibrotic role of TGFβ1 in adipose tissue characterized by collagen accumulation, hypoxia (which upregulates VEGFA [79]), and M1 macrophage polarization [26, 79–83].

### 4.5 Age-related differences

In our dataset, aging was correlated with a decrease in the number of adipocytes (**Figure 8A**, - 0.549) and percentage of lipids (**Figure 8B**, −0.601). It is important to note that age and adipocyte size had no correlation (−0.078), indicating that the observed decrease in the number of adipocytes with age was not due to an increase in the size (hypertrophy) of adipocytes. Instead, our findings are consistent with other studies that have shown hyperplastic growth in adipose tissue declines with age [21]. Furthermore, the decrease in lipid percent with age, and the lack of correlation between BMI and percentage of lipids (−0.186) in our subcutaneous adipose tissue samples, supports studies that show a shift in lipid storage from subcutaneous to visceral depots with age [65–66]. The ability to buffer lipids in adipocytes declines with age, leading to ectopic lipid deposition in the liver and muscle [84]. Supporting the shift in adipocyte lipid storage capacity, there was an increase in the number of extracellular lipid droplets with age (**Figure 8D**, 0.370).

**Figure 8.**
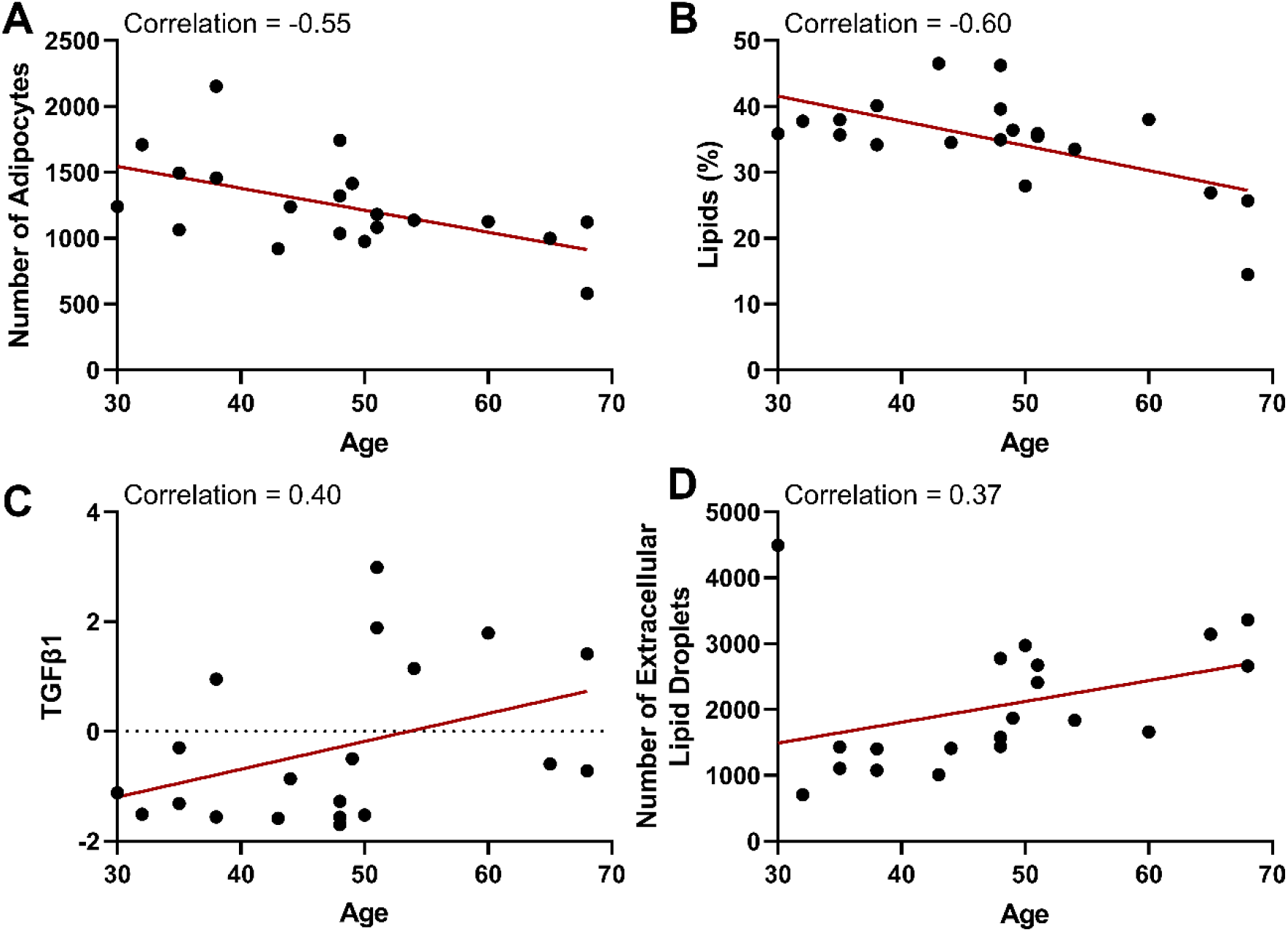
With increasing age there is a decline in the number of adipocytes and the percentage of lipids and an increase in TGFβ1 and the number of lipid droplets deposited extracellularly. The age of patients at the time of surgery is plotted versus the number of adipocytes measured in histological images (A), the lipid percent measured in the same images (B), the gene expression of TGFβ1 (C), and the number of extracellular lipid droplets counted in histological images (D). TGFβ1 gene expression is represented as a delta CT from the housekeeping gene (ΔCT = CT_SDHA_ – CT_Target gene_). With the formula used, gene expression is relative to the housekeeping gene, and increases from a negative value to a higher positive value. Correlations from the correlation matrix are indicated on each plot.

As mentioned previously, aging was correlated with an increase in TGFβ1 gene expression (**Figure 8C**, 0.40) and upregulation of a M1 macrophage marker (CD86, 0.33), while displaying no correlation with a M2 macrophage marker (CD163, 0.05). It is known that aging results in compounded Reactive Oxidative Species which readily primes macrophages toward a proinflammatory M1 state and promotes age-related metabolic syndrome (atherosclerosis, obesity, and type II diabetes) [85]. TGFβ signaling largely depends on the cell type and context. However, the upregulation of TGFβ ligands with age is known to contribute to cell degeneration, tissue fibrosis, inflammation, decreased regeneration capacity, and metabolic malfunction in other tissues [86]. Therefore, age-associated changes in subcutaneous adipose tissue are an important context that warrants further investigation.

### 4.6 Ancestral differences

Race was the highest loading in the fourth dimension, which accounted for over 11% of the variance in the dataset. Despite known differences in disease prevalence in diverse patient populations, cellular ancestry is largely overlooked in biomedical research [87]. This ancestry-blind approach ignores important differences in disease progression that are key to better therapeutic options; such as ancestral differences in cellular transcription [88] and drug responsiveness [89]. Adipose tissue plays a central role in obesity and type II diabetes progression with many studies indicate a greater prevalence in Hispanic and Black patients over White patients [90–92].

In the current study, samples derived from self-identified Black patients had significantly lower gene expression levels of TGFβ1 and ACTA2 (**Figure 9A,B**) and a trend towards lower hydroxyproline collagen content (**Figure 9C**) than White patients (all patients self-identified as non-Hispanic). As mentioned previously, fibrosis is characterized by enhanced collagen deposition, driven by high levels of TGFβ1 and its target gene ACTA2 that induces a myofibroblastic phenotype [26, 77]. TGFβ signals resolution of inflammatory signals, and therefore attenuated TGFβ/Smad signaling results in a reduced mechanism to resolve inflammation [93]. The observed low fibrotic signature in adipose tissues samples derived from Black patients is consistent with Black patients having lower incidences of pulmonary fibrosis than non-Hispanic White patients [94]. Therefore, future work should explore the role of ancestry in subcutaneous adipose tissue fibrosis.

**Figure 9.**
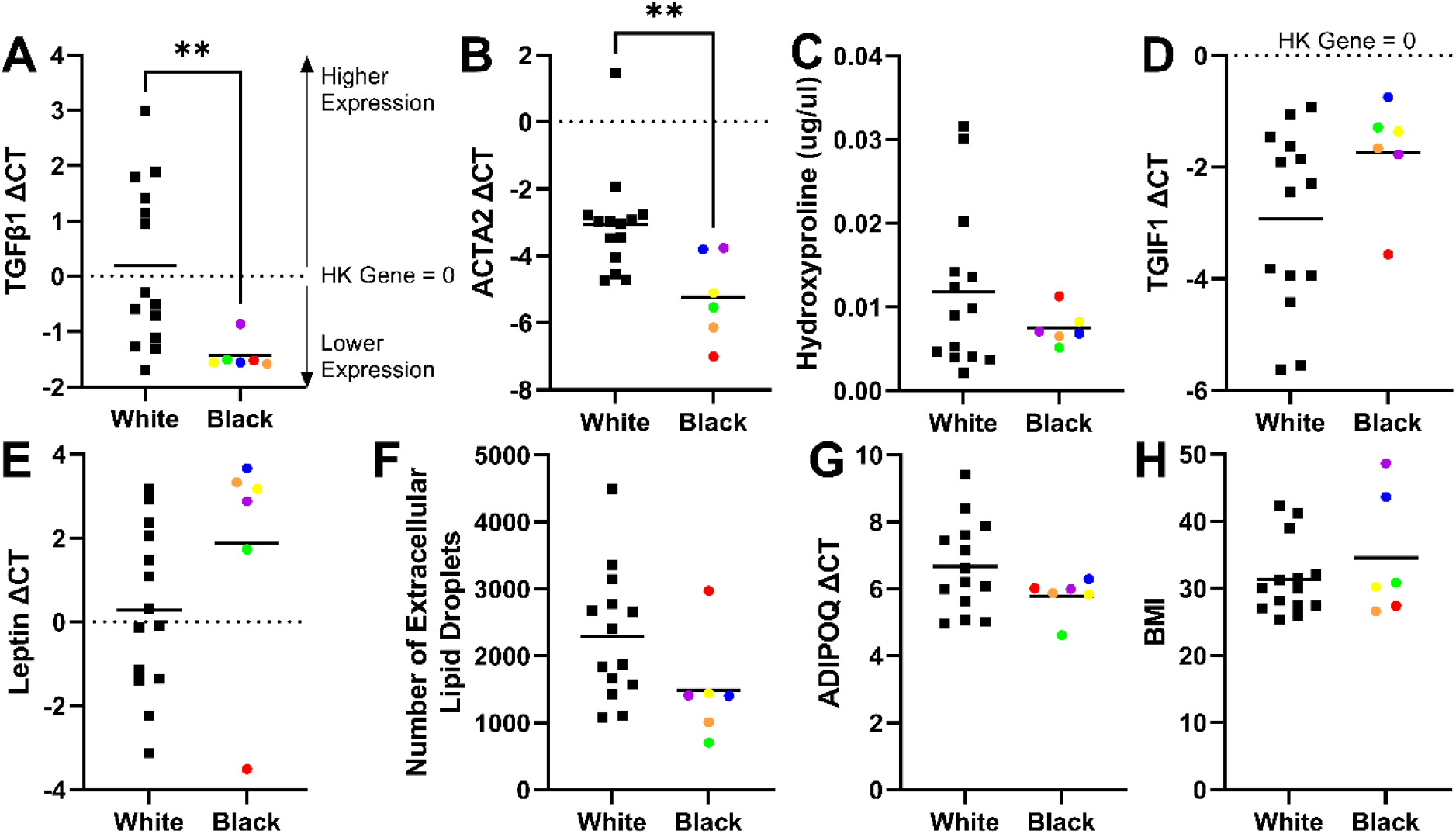
Significantly lower fibrotic gene expression (TGFβ1 and ACTA2) and trends related to collagen content, TGIF1, leptin, extracellular lipid droplets, and adiponectin were observed in Black patients compared to White patients and were independent of BMI. Samples derived from self-identified Black patients had significantly lower gene expression levels of TGFβ1 (A) and ACTA2 (B) and a trend towards lower hydroxyproline collagen content (C) than White patients (all patients identified as non-Hispanic). On average, samples from Black patients also had higher gene expression levels of TGIF1 (D) and leptin (E), and fewer extracellular lipid droplets (F) compared to White patients. Adiponectin levels were also lower in Black patient samples compared to White (G). Differences between the population groups are independent of body mass index (BMI), as there was no significant differences in BMI between the groups (H). Samples derived from Black patients were color coded by patient source to highlight that the outliers in TGIF1, leptin, hydroxyproline, and number of extracellular lipid droplets were all from the same patient (red dots). This sample follows the same trends as the aggregated data (with combined ancestries) that reduced levels of TGIF1 are linked with higher numbers of extracellular lipid droplets and an inability to suppress fibrotic changes in adipose tissue. Statistical significance was determined by an unpaired t-test where ** indicates p<0.01.

Our data supports the concept that disease progression could present differently in different patient populations. For example, there is an observed trend that Hispanic patients disproportionally develop Nonalcoholic fatty liver disease (NAFLD) and have high rates of obesity, visceral adipose tissue, and insulin resistance. In contrast, Black patients tend to have a high prevalence of obesity and insulin resistance with a paradoxically favorable lipid profile and low prevalence of visceral adipose tissue and NAFLD [95]. On average, samples from Black patients in our study had higher gene expression levels of TGIF1 (**Figure 9D**) and leptin (**Figure 9E**) with fewer extracellular lipid droplets (**Figure 9F**) compared to samples from White patients. We speculate there could be a metabolic phenotype in some patients that favors intracellular lipid accumulation with high TGIF1 transcription that limits extracellular lipid release and (at least initially) favors a diabetic disease process over disease processes that are driven by lipid accumulation in other tissues. This is consistent with a recent study indicating that plasma triglyceride concentrations are lower in African Americans compared to non-Hispanic White, Hispanic White, East Asian, and South Asian ethnicities (controlling for the level of insulin sensitivity) [96]. Furthermore, preliminary findings from another research group indicated that TGIF1 overexpression leads to severe hyperglycemia and obesity-associated diabetes [97], suggesting a link between high TGIF1 signaling and an increased likelihood of developing a diabetic phenotype. The FAMD analysis also identified an interaction between sample ancestry and adiponectin gene expression. While not significant (by an unpaired t-test), lower levels of adiponectin were observed in samples derived from Black patients. Adiponectin is an insulin-sensitizing, anti-inflammatory, and anti-diabetic adipokine that’s levels decrease with increasing adipose tissue mass [98–100]. Therefore, lower adiponectin levels (with similar BMI ranges, **Figure 9H**) further support the higher observed rates of insulin resistance in Black patients compared to White patients [90–92]. Therefore, it is imperative that future work explores TGIF1 signaling, lipid release mechanisms from adipocytes, diabetic disease progression, and mechanisms of inflammation resolution in cells derived from diverse backgrounds.

Interestingly, the outliers in TGIF1, leptin, hydroxyproline, and number of extracellular lipid droplets were all the same patient (red dots). This further supports the aggregated data (with combined ancestries) that reduced levels of TGIF1 are linked with higher numbers of extracellular lipid droplets and an inability to suppress fibrotic changes in adipose tissue. It should also be noted that our sample size is small for this study, and we had uneven sample sizes for each group, with only 6 samples from self-identified Black patients and 14 from self-identified White patients. However, even with the small sample size from Black patients the results are independent of BMI with a similar spread of BMIs between the groups (**Figure 9H**). Collectively, these results underscore the urgent need to account for patient ancestry in biomedical research.

### 4.7 Macrophage surface markers

Macrophages play a major role in tissue homeostasis and disease, modulating the growth of tissues as well as tissue remodeling and organization. As mentioned previously, CD86 is considered a classically activated M1 macrophage marker and CD163 is an alternatively activated M2 macrophage marker [70–71] which were positively correlated in our dataset (**Figure 10A**). While macrophage polarization results in distinct functional phenotypes that are often thought of as divergent (i.e.; an increase in M1 polarization is associated with a decrease in M2 polarization), induction routes are complex, interconnecting network systems rather than simple switches in activation patterns [101]. It is well established that lean adipose tissue contains resident macrophages that are alternatively activated (M2) and maintain tissue homeostasis [89]. As adipose tissue increases in mass, there is infiltration of additional macrophages and the number of classically activated (M1) macrophages increases contributing to the inflammatory signature of obesity and insulin resistance [102–103], where there is an observed phenotypic switch from one activation state to another [104–105]. However, there is growing evidence that macrophage activation patterns within adipose tissue are more complicated than a full switch from M2 to M1 in obesity, which could be related to significant differences in macrophage profiles in animals and human models [106]. As mentioned previously, in humans, high levels of both CD86 and CD163 gene expression have been identified in samples from overweight/obese patients [71]. Consistent with this finding, both CD86 and CD163 also showed a high correlation with leptin in our dataset (0.60 and 0.64, respectively), a hormone that is directly related to total body fat mass in humans [43]. Therefore, our results indicate that higher levels of both M1 and M2 polarization markers are present in human subcutaneous adipose tissue with increasing body fat mass.

**Figure 10.**
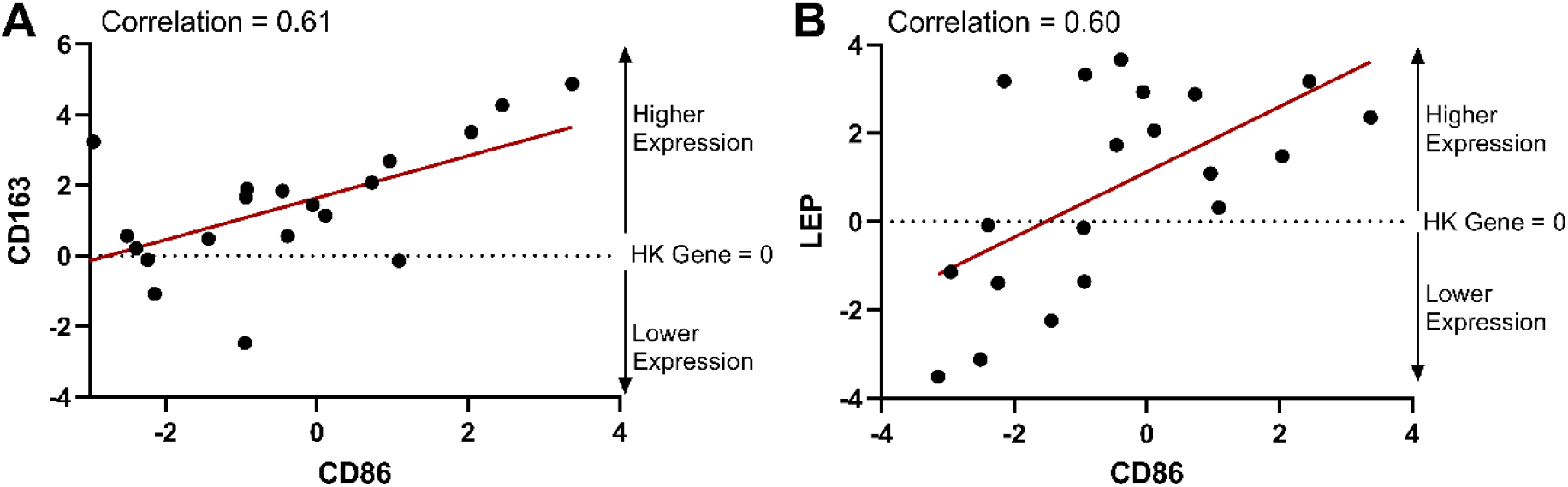
Gene expression of the clasically activated CD86 macrophage surface marker and the alternatively activated CD163 surface marker were correlated with each other and with leptin gene expression. CD86 gene expression is plotted versus CD163 gene expression (A) and leptin (LEP) gene expression (B). Gene expression is represented as a delta CT from the housekeeping (HK) gene (ΔCT = CT_SDHA_ – CT_Target gene_). Therefore, the HK gene is equal to 0 on the plots. With the formula used, gene expression is relative to the housekeeping gene, and increases from a negative value to a higher positive value. Correlations from the correlation matrix are indicated on each plot.

Other correlations that were consistent for both macrophage surface markers, CD86 and CD163, for example: adipocyte diameter (0.46 and 0.33, respectively) and TGIF1 (mentioned previously and shown in **Figure 6**). Hypertrophic adipocytes are known to become inflammatory and necrotic, attracting M1 macrophages, which organize into “crown-like” structures that eventually clear necrotic adipocytes [107–109]. M2 macrophages regulate adipose progenitors by initiating proliferation and differentiation into adipocytes to mitigate existing adipocytes’ nutritional overload and prevent their further enlargement [110]. Therefore, it is unsurprising that both cell types correlate with hypertrophic adipocytes.

While the macrophage surface markers were correlated in our dataset, there were also distinct correlations associated with each polarization phenotype. For example, higher M1 CD86 gene expression was correlated with TGFβ1 (0.46 versus 0 for CD163), increasing age (0.33 versus 0.05 for CD163), and metabolic activity (−0.41 versus 0.14 for CD163) as discussed previously. While higher M2 CD163 gene expression was correlated with a higher overall lipid percent (0.45 versus 0.18 for CD86), which is likely related to the homeostatic role of M2 macrophages in recruiting the differentiation of new adipocytes [110].

### 4.8 Diabetic status and AGEs

The largest loading in dimensions 6 and 7 of the FAMD was diabetic status (**Table 2**, combined this accounts for ∼12% of the variance). Diabetic status was clustered with AGEs and DNA content for dimensions 6 and 7, respectively (**Figure 5**). The largest loading in the final dimension of the FAMD (that was included to account for >80% of the variance) was AGEs and was clustered with diabetic status in that dimension as well. The AGE content in the tissue samples varied considerably (**Supplemental Figure 3F**). However, patients with diabetes or that formerly had diabetes had higher levels of AGEs. Patients with a higher BMI also had higher levels of AGEs. This trend is supported by literature that shows increased AGE content in tissue is associated with age, diabetes, and obesity-related complications [111–112].

### 4.9 Smoking Status

The patients’ smoking status was the largest loading in the eighth dimension of the FAMD (**Table 2, Figure 5**). Adipose tissue has nicotinic receptors [113] that enable tobacco to greatly affect adipose tissue properties and cause downstream effects. The FAMD indicated that diabetic status and smoking were clustered. Diabetic patients who smoke have health problems, comorbidities, and poor outcomes compared to diabetic patients that do not smoke [114–117].

### 4.10 Limitations

The correlative nature of our investigation limits the conclusions that can be drawn in terms of cause and effect. Instead, we view this study as complementing current literature and informing future research directions into cause-and-effect relationships. It is also important to acknowledge the parameters of this study. The tissue samples used in this study were all from patients that were either overweight or obese (BMI>25). Therefore, the correlations drawn from the data are not applicable to lean patients. Furthermore, tissue was taken from the subcutaneous adipose depot, and therefore are not applicable to other adipose depots in the body. All gene expression values are also taken from the bulk tissue; therefore cell-type specific gene expression is not possible (for example an upregulation of ACTA2 could mean there are more myofibroblasts or adipocytes are increasing their expression of the gene). As is the case for most studies, this preliminary study of 20 patient samples would be strengthened by the addition of more patients and an increase in the diversity of the patient population. For example, no patient samples were collected from Hispanic backgrounds. Therefore, replicating this study in locations with access to different patient demographics is essential. Additionally, more information on comorbidities, genetics, and other lifestyle information such as history of exercise, diet, and smoking duration and cessation would strengthen and expand these results providing valuable information for disease treatment and modeling.

## 5. Conclusions

Many of the results in this study agree with well-established findings in animal studies and other *in vitro* model systems. For example, TGFβ1 was associated with adipose tissue fibrosis, which is characterized by collagen accumulation and M1 macrophage polarization [26, 80–83] and diabetes was associated with AGE [111–112]. This study also revealed new patterns of key markers that drive human variability in adipose tissues.

VEGFA and ACTA2 gene expression were the highest loadings in the first two dimensions of the FAMD, respectively. Vascularization of human adipose tissue varies considerably between patients, and in this study VEGFA was correlated with adiponectin gene expression. ACTA2 induces a so-called “cellular identity crisis” as described elsewhere [76], which we found was correlated with higher collagen content and TGFβ1 signaling, a reduction in adipocyte cellular diameter, and an upregulation in VEGFA gene expression. There was also a key role of TGIF1 in accounting for variability in the adipose tissues, which is only recently being uncovered. This study emphasizes the importance of TGIF1 signaling for driving patient differences with a strong correlation to leptin signaling.

The number of adipocytes was the highest loading in the third dimension of the FAMD, where a decrease in the density of adipocytes was associated with aging and an increase in cellular proliferative capacity of ASCs. Aging was also associated with a decrease in overall lipid percentage that favored lipid deposition extracellularly, an increase in TGFβ1, and an increase in M1 macrophage polarization.

The highest loading in the fourth dimension of the FAMD, which accounted for over 11% of the variability in patient-related differences in this study, was self-identified race. While our sample size was small, Black patients had significantly lower gene expression levels of TGFβ1 and ACTA2. This finding indicates there is an urgent need to account for patient ancestry in biomedical research to develop better therapeutic strategies for all patients.

Amongst many other findings, this study revealed an interesting gene expression pattern where M1 and M2 macrophage markers were correlated with each other and leptin (for BMI >25). This finding supports growing evidence that macrophage polarization in obesity involves a complex, interconnecting network system rather than a simple switch in activation patterns from M2 to M1 with increasing body mass. Therefore, the characterization of M2 macrophages in overweight/obese tissues warrants further investigation.

## Acknowledgements

This work was funded (in part) by the Dowd Fellowship from the College of Engineering at Carnegie Mellon University. The authors would like to thank Philip and Marsha Dowd for their financial support and encouragement. This work was also funded (in part) through the NIH-T32 Biomechanics and Regenerative Medicine (BiRM) Training Grant. The authors would like to acknowledge the Department of Plastic Surgery at the University of Pittsburgh’s Medical Center and the Adipose Stem Cell Center for procuring tissue samples used in this work. The authors would also like to thank Mallory Griffin for help in isolating tissues and Isabelle Chickanosky and Maya Garg for assisting in gathering doubling time data.

## Supplementary Materials

### Supplement 1. Statistical Significance for Image Analysis

As would be expected given the diversity of the human samples and how adipocytes vary considerably in size from 20-300 μm [3], there were statistical differences in adipocyte diameters between patients (**Supplemental Figure 1**). The average adipocyte diameter in this study ranged from 55.05 μm (Patient 10) to 88.89 μm (Patient 12) which is within the normal range for human subcutaneous adipose tissue. 6/20 patients were classified as “large” adipocytes as their diameter was on average >70 μm [118]. It is known that adipocyte size affects adipocyte function, where large (70-120 μm) and very large (>120 μm) diameter adipocytes have reduced insulin sensitivity, lower metabolic health, and increased proinflammatory cytokine and free fatty acid secretion [118–119].

The number of adipocytes in each sample was the highest loading in the third dimension of the FAMD (**Table 2**), indicating this was a key driver of variability in human subcutaneous adipose tissues. The correlation matrix indicated that the number of adipocytes decreased with increasing age (−0.55) and as the diameter of the adipocyte increased (−0.47) (**Supplemental Figure 2**). It should be noted that images were taken at the same magnification, therefore large adipocytes would result in fewer cells per frame. However, not all images contained densely packed adipocytes, with many containing collagen-dense areas and extracellular lipid droplets (**Figure 2**). Another interesting finding was the correlation between the number of adipocytes and a longer doubling time of the stromal vascular fraction (**Supplemental Figure 2B**). The more adipocytes there were, the longer it took the stromal vascular fraction to proliferate, indicating hyperplastic growth in the organ limited the cellular proliferative capacity of the remaining preadipocyte population.

The total number of extracellular lipid droplets were counted for each patient (**Supplemental Figure 1J**). Most patients had significantly more extracellular lipid droplets than adipocytes, except for patients 9, 11, 18, and 19. Interestingly, even though there is variability in the number of extracellular lipid droplets to adipocytes, there are no statistically significant differences in the co-localization score of the number of extracellular lipid droplets in collagen/total number of extracellular lipid droplets (**Supplemental Figure 1K**). The co-localization scores are all greater than 0.5, indicating that the majority of extracellular lipid droplets detected were found in collagen-dense areas.

**Supplemental Figure 1.**
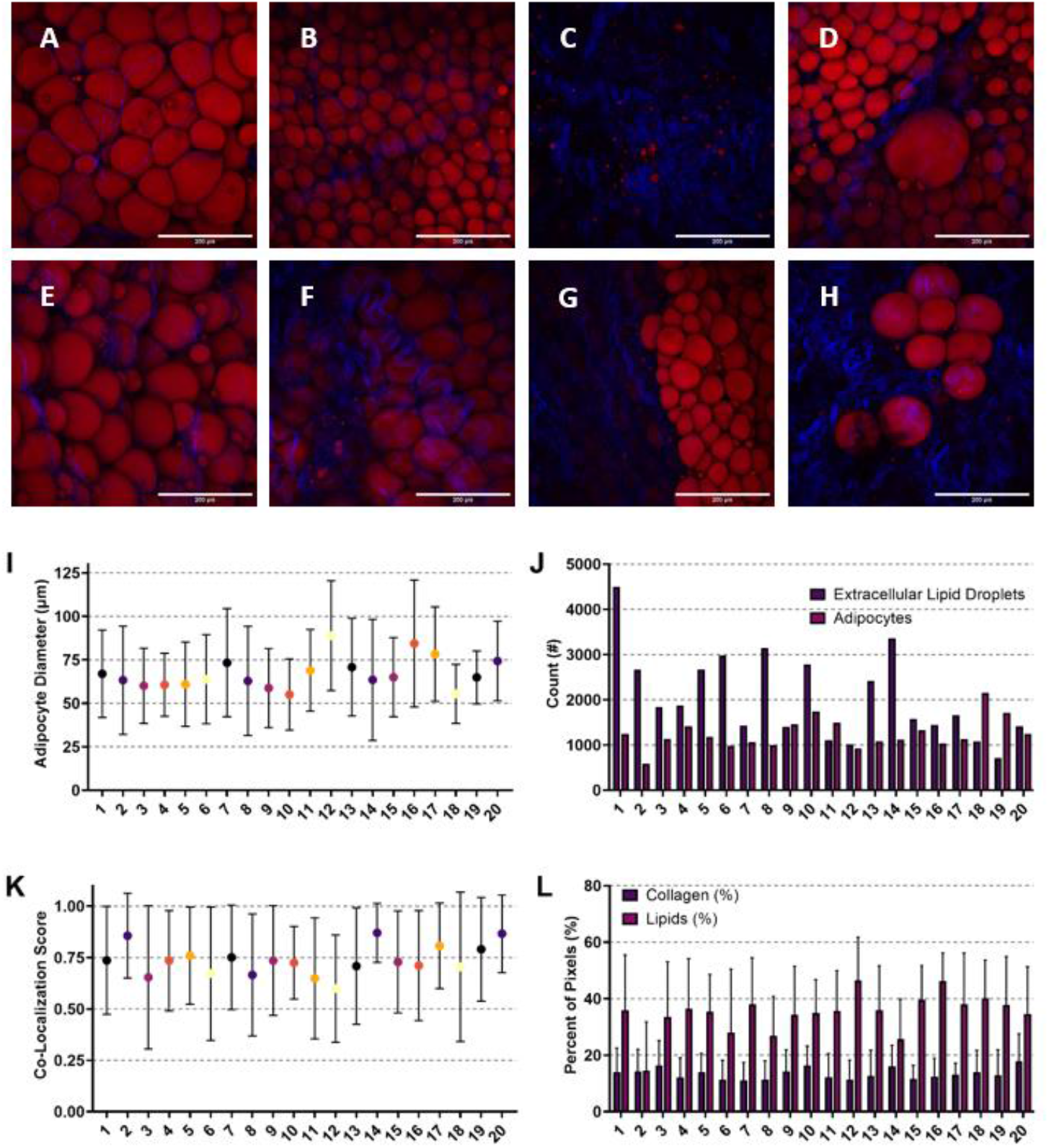
While there is considerable variability in size of adipocytes, number of adipocytes, and percentage of lipid/collagen, extracellular lipid droplets are consistently found in collagen-dense regions. Representative images used to analyze number of adipocytes, adipocyte size, extracellular lipid location, and the percentage of lipids and collagen (A-H). Images show lipid droplets (red) and collagen (blue). Scale bars are 200 μm. Each image is from a different patient. Patients had variable adipocyte diameters (I), quantity of extracellular lipid droplets and adipocytes (J), and quantity of collagen (L). However, there was no statistical difference in the co-localization score (K) defined as the number of extracellular lipid droplets in collagen/total number of extracellular lipid droplets. High values for the colocalization score signify more extracellular lipid droplets were found in collagenous regions than in non-collagenous regions of the tissue. This indicates that large variations in the quantity of extracellular lipid droplets and their ratio to adipocytes did not significantly change the location of where the extracellular lipid droplets were distributed (primarily in collagenous regions).

**Supplemental Figure 2.**
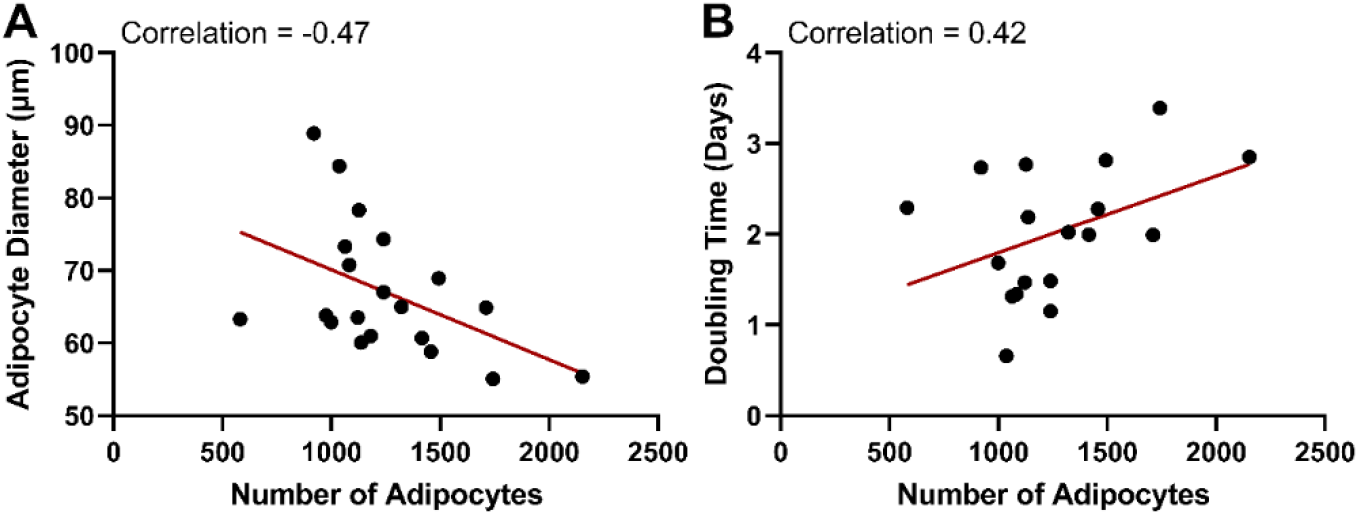
As the number of adipocytes increases the diameter of the remaining adipocytes decreases and the doubling time of the stromal vascular fraction gets longer. The number of adipocytes in histological images was plotted versus the diameter of adipocytes in the same images (A) and against the recorded doubling time of the stromal vascular fraction seeded into cell culture flasks. Correlations from the correlation matrix are indicated on each plot.

**Supplemental Table 1.**
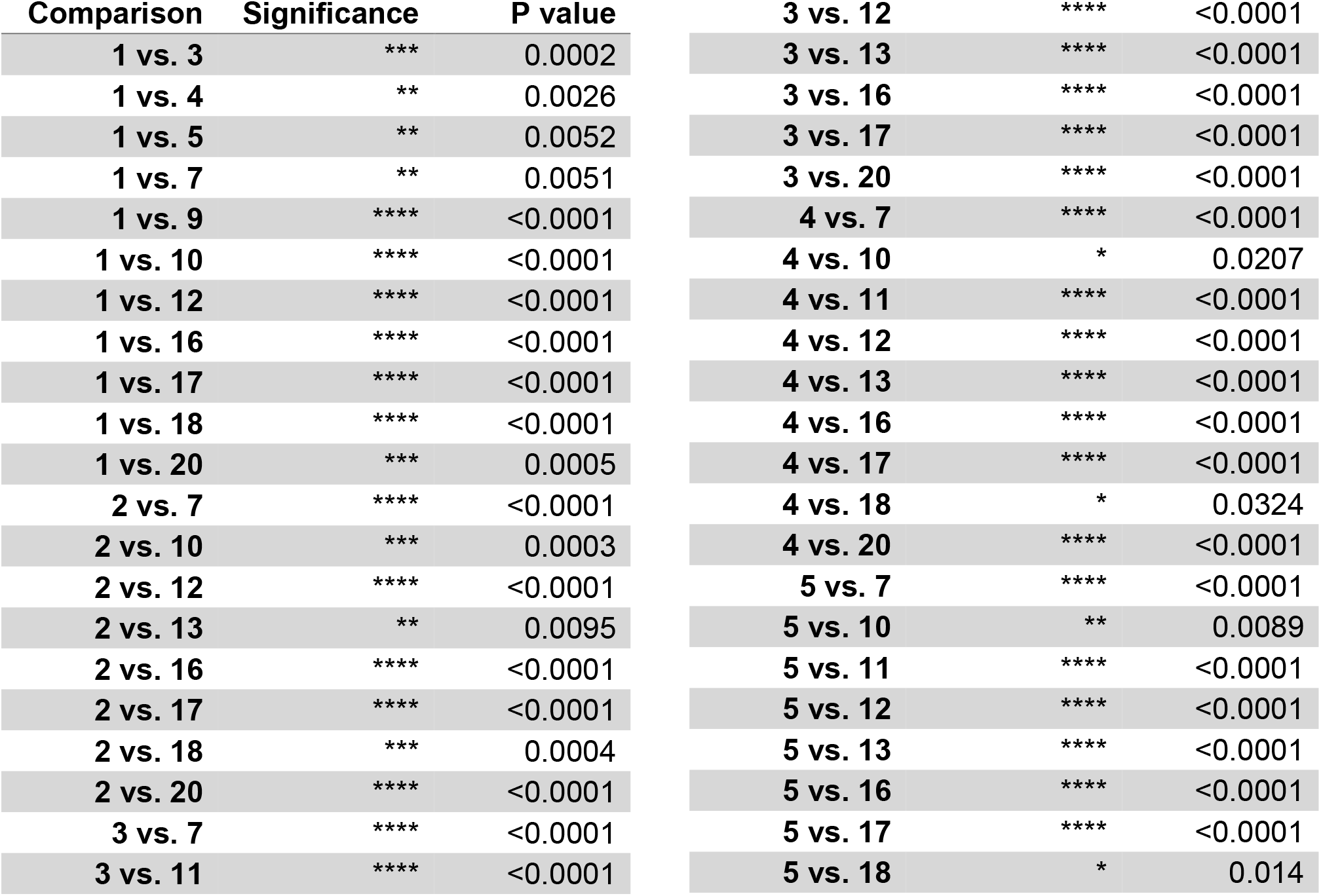
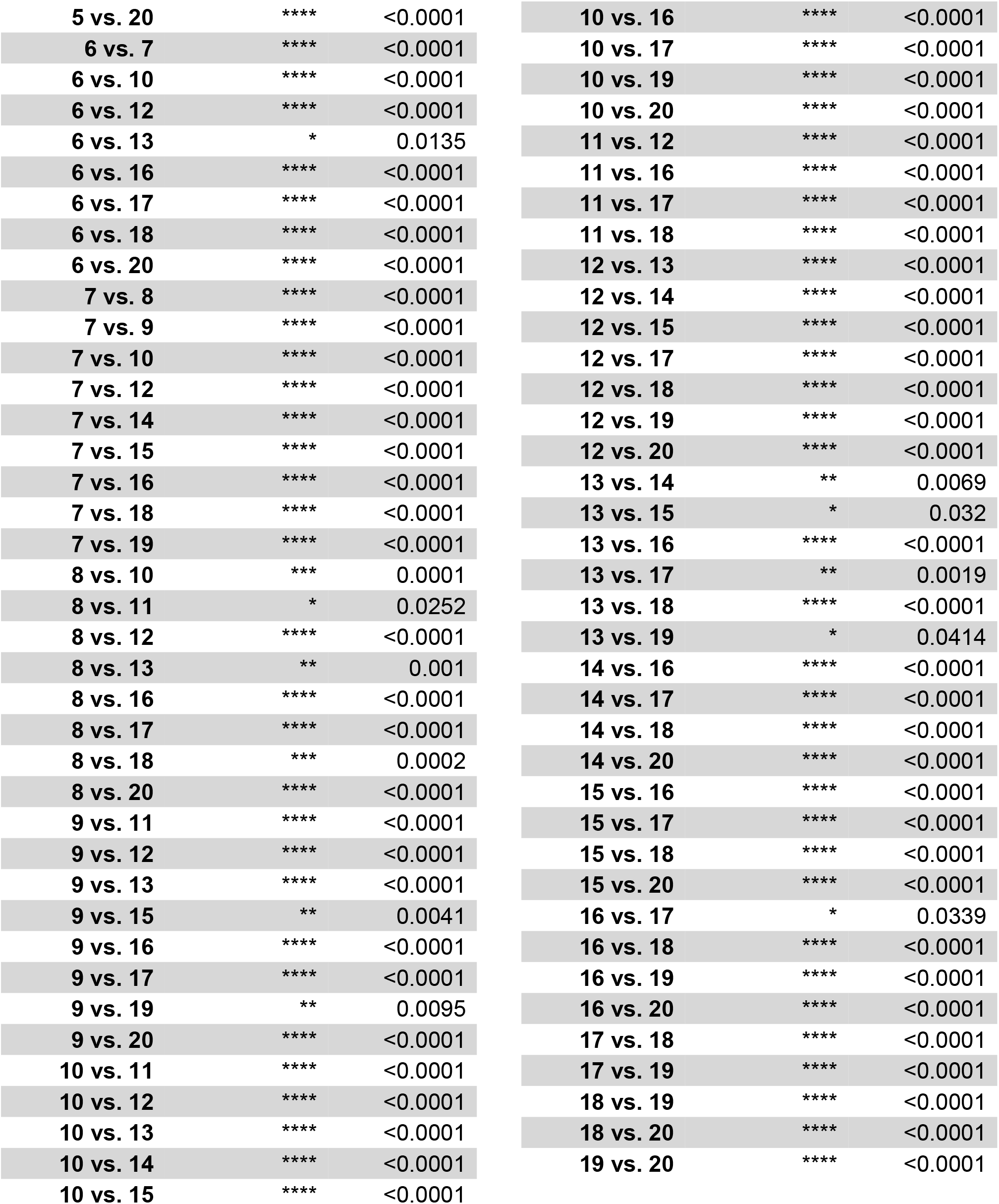
Statistical significance for adipocyte size. Determined through a one-way ANOVA followed by Tukey’s post-hoc analysis. Non-statistically significant comparisons are not shown. Significance was defined as p<0.05. * p<0.05, ** p<0.01, *** p<0.005, **** p<0.0001.

| Comparison | Significance | P value |  |  |  |
| --- | --- | --- | --- | --- | --- |
| 1 vs. 3 | *** | 0.0002 | 3 vs. 12 | **** | <0.0001 |
| 1 vs. 4 | ** | 0.0026 | 3 vs. 13 | **** | <0.0001 |
| 1 vs. 5 | ** | 0.0052 | 3 vs. 16 | **** | <0.0001 |
| 1 vs. 7 | ** | 0.0051 | 3 vs. 17 | **** | <0.0001 |
| 1 vs. 9 | **** | <0.0001 | 3 vs. 20 | **** | <0.0001 |
| 1 vs. 10 | **** | <0.0001 | 4 vs. 7 | **** | <0.0001 |
| 1 vs. 12 | **** | <0.0001 | 4 vs. 10 | * | 0.0207 |
| 1 vs. 16 | **** | <0.0001 | 4 vs. 11 | **** | <0.0001 |
| 1 vs. 17 | **** | <0.0001 | 4 vs. 12 | **** | <0.0001 |
| 1 vs. 18 | **** | <0.0001 | 4 vs. 13 | **** | <0.0001 |
| 1 vs. 20 | *** | 0.0005 | 4 vs. 16 | **** | <0.0001 |
| 2 vs. 7 | **** | <0.0001 | 4 vs. 17 | **** | <0.0001 |
| 2 vs. 10 | *** | 0.0003 | 4 vs. 18 | * | 0.0324 |
| 2 vs. 12 | **** | <0.0001 | 4 vs. 20 | **** | <0.0001 |
| 2 vs. 13 | ** | 0.0095 | 5 vs. 7 | **** | <0.0001 |
| 2 vs. 16 | **** | <0.0001 | 5 vs. 10 | ** | 0.0089 |
| 2 vs. 17 | **** | <0.0001 | 5 vs. 11 | **** | <0.0001 |
| 2 vs. 18 | *** | 0.0004 | 5 vs. 12 | **** | <0.0001 |
| 2 vs. 20 | **** | <0.0001 | 5 vs. 13 | **** | <0.0001 |
| 3 vs. 7 | **** | <0.0001 | 5 vs. 16 | **** | <0.0001 |
| 3 vs. 11 | **** | <0.0001 | 5 vs. 17 | **** | <0.0001 |
|  |  |  | 5 vs. 18 | * | 0.014 |

|  |  |  |
| --- | --- | --- |
| 5 vs. 20 | **** | <0.0001 |
| 6 vs. 7 | **** | <0.0001 |
| 6 vs. 10 | **** | <0.0001 |
| 6 vs. 12 | **** | <0.0001 |
| 6 vs. 13 | * | 0.0135 |
| 6 vs. 16 | **** | <0.0001 |
| 6 vs. 17 | **** | <0.0001 |
| 6 vs. 18 | **** | <0.0001 |
| 6 vs. 20 | **** | <0.0001 |
| 7 vs. 8 | **** | <0.0001 |
| 7 vs. 9 | **** | <0.0001 |
| 7 vs. 10 | **** | <0.0001 |
| 7 vs. 12 | **** | <0.0001 |
| 7 vs. 14 | **** | <0.0001 |
| 7 vs. 15 | **** | <0.0001 |
| 7 vs. 16 | **** | <0.0001 |
| 7 vs. 18 | **** | <0.0001 |
| 7 vs. 19 | **** | <0.0001 |
| 8 vs. 10 | *** | 0.0001 |
| 8 vs. 11 | * | 0.0252 |
| 8 vs. 12 | **** | <0.0001 |
| 8 vs. 13 | ** | 0.001 |
| 8 vs. 16 | **** | <0.0001 |
| 8 vs. 17 | **** | <0.0001 |
| 8 vs. 18 | *** | 0.0002 |
| 8 vs. 20 | **** | <0.0001 |
| 9 vs. 11 | **** | <0.0001 |
| 9 vs. 12 | **** | <0.0001 |
| 9 vs. 13 | **** | <0.0001 |
| 9 vs. 15 | ** | 0.0041 |
| 9 vs. 16 | **** | <0.0001 |
| 9 vs. 17 | **** | <0.0001 |
| 9 vs. 19 | ** | 0.0095 |
| 9 vs. 20 | **** | <0.0001 |
| 10 vs. 11 | **** | <0.0001 |
| 10 vs. 12 | **** | <0.0001 |
| 10 vs. 13 | **** | <0.0001 |
| 10 vs. 14 | **** | <0.0001 |
| 10 vs. 15 | **** | <0.0001 |
| 10 vs. 16 | **** | <0.0001 |
| 10 vs. 17 | **** | <0.0001 |
| 10 vs. 19 | **** | <0.0001 |
| 10 vs. 20 | **** | <0.0001 |
| 11 vs. 12 | **** | <0.0001 |
| 11 vs. 16 | **** | <0.0001 |
| 11 vs. 17 | **** | <0.0001 |
| 11 vs. 18 | **** | <0.0001 |
| 12 vs. 13 | **** | <0.0001 |
| 12 vs. 14 | **** | <0.0001 |
| 12 vs. 15 | **** | <0.0001 |
| 12 vs. 17 | **** | <0.0001 |
| 12 vs. 18 | **** | <0.0001 |
| 12 vs. 19 | **** | <0.0001 |
| 12 vs. 20 | **** | <0.0001 |
| 13 vs. 14 | ** | 0.0069 |
| 13 vs. 15 | * | 0.032 |
| 13 vs. 16 | **** | <0.0001 |
| 13 vs. 17 | ** | 0.0019 |
| 13 vs. 18 | **** | <0.0001 |
| 13 vs. 19 | * | 0.0414 |
| 14 vs. 16 | **** | <0.0001 |
| 14 vs. 17 | **** | <0.0001 |
| 14 vs. 18 | **** | <0.0001 |
| 14 vs. 20 | **** | <0.0001 |
| 15 vs. 16 | **** | <0.0001 |
| 15 vs. 17 | **** | <0.0001 |
| 15 vs. 18 | **** | <0.0001 |
| 15 vs. 20 | **** | <0.0001 |
| 16 vs. 17 | * | 0.0339 |
| 16 vs. 18 | **** | <0.0001 |
| 16 vs. 19 | **** | <0.0001 |
| 16 vs. 20 | **** | <0.0001 |
| 17 vs. 18 | **** | <0.0001 |
| 17 vs. 19 | **** | <0.0001 |
| 18 vs. 19 | **** | <0.0001 |
| 18 vs. 20 | **** | <0.0001 |
| 19 vs. 20 | **** | <0.0001 |

**Supplemental Table 2.** Statistical significance for collagen and lipid pixel percent. Determined through a two-way ANOVA followed by Tukey’s post-hoc analysis. Non-statistically significant comparisons are not shown. Significance was defined as p<0.05. * p<0.05, ** p<0.01, *** p<0.005, **** p<0.0001.

| Comparison | Significance | P value |
| --- | --- | --- |
| 1: Collagen (%) vs. 1: Lipids (%) | *** | 0.0002 |
| 1: Lipids (%) vs. 2: Lipids (%) | *** | 0.0003 |
| 2: Lipids (%) vs. 3: Lipids (%) | ** | 0.0040 |
| 2: Lipids (%) vs. 4: Lipids (%) | *** | 0.0002 |
| 2: Lipids (%) vs. 5: Lipids (%) | *** | 0.0005 |
| 2: Lipids (%) vs. 7: Lipids (%) | **** | <0.0001 |
| 2: Lipids (%) vs. 9: Lipids (%) | ** | 0.0020 |
| 2: Lipids (%) vs. 10: Lipids (%) | *** | 0.0009 |
| 2: Lipids (%) vs. 11: Lipids (%) | *** | 0.0004 |
| 2: Lipids (%) vs. 12: Lipids (%) | **** | <0.0001 |
| 2: Lipids (%) vs. 13: Lipids (%) | *** | 0.0003 |
| 2: Lipids (%) vs. 15: Lipids (%) | **** | <0.0001 |
| 2: Lipids (%) vs. 16: Lipids (%) | **** | <0.0001 |
| 2: Lipids (%) vs. 17: Lipids (%) | **** | <0.0001 |
| 2: Lipids (%) vs. 18: Lipids (%) | **** | <0.0001 |
| 2: Lipids (%) vs. 19: Lipids (%) | **** | <0.0001 |
| 2: Lipids (%) vs. 20: Lipids (%) | ** | 0.0014 |
| 3: Collagen (%) vs. 3: Lipids (%) | * | 0.0234 |
| 4: Collagen (%) vs. 4: Lipids (%) | **** | <0.0001 |
| 5: Collagen (%) vs. 5: Lipids (%) | *** | 0.0003 |
| 6: Collagen (%) vs. 6: Lipids (%) | * | 0.0385 |
| 6: Lipids (%) vs. 12: Lipids (%) | ** | 0.0062 |
| 6: Lipids (%) vs. 16: Lipids (%) | ** | 0.0086 |
| 7: Collagen (%) vs. 7: Lipids (%) | **** | <0.0001 |
| 8: Lipids (%) vs. 12: Lipids (%) | ** | 0.0021 |
| 8: Lipids (%) vs. 16: Lipids (%) | ** | 0.0030 |
| 9: Collagen (%) vs. 9: Lipids (%) | ** | 0.0014 |
| 10: Collagen (%) vs. 10: Lipids (%) | ** | 0.0061 |
| 11: Collagen (%) vs. 11: Lipids (%) | **** | <0.0001 |
| 12: Lipids (%) vs. 14: Lipids (%) | *** | 0.0005 |
| 14: Lipids (%) vs. 16: Lipids (%) | *** | 0.0008 |
| 15: Collagen (%) vs. 15: Lipids (%) | **** | <0.0001 |
| 16: Collagen (%) vs. 16: Lipids (%) | **** | <0.0001 |
| 17: Collagen (%) vs. 17: Lipids (%) | **** | <0.0001 |
| 18: Collagen (%) vs. 18: Lipids (%) | **** | <0.0001 |
| 19: Collagen (%) vs. 19: Lipids (%) | **** | <0.0001 |
| 20: Collagen (%) vs. 20: Lipids (%) | * | 0.0359 |

### Supplement 2. Mechanical properties, DNA content, collagen content, AGEs, metabolic activity, and doubling rate

The elastic modulus, peak stress, stromal vascular fraction (SVF) doubling time, hydroxyproline content, metabolic activity, and advanced glycation end-products (AGEs) content are shown for each patient (**Supplemental Figure 3**). There is large variability between all of the samples for each metric, therefore statistical significance between patients is not shown on graphs and is represented in tabular form (**Supplemental Table 3, Supplemental Table 4, Supplemental Table 5, Supplemental Table 6**).

As indicated on the correlation matrix (**Figure 3**), the highest correlation between quantitative variables in this study (correlation = 0.94) was between the Elastic Modulus and the Peak Stress (**Supplemental Figure 4**). Because these two variables are both gathered from compressive testing, the high correlation is not surprising. Elastic Modulus and Peak Stress define different material behaviors, i.e. the ability of the material to resist elastic deformation (“stiffness” defined as the slope of the stress/strain curve in the elastic region) and the maximum stress (strength) the material withstands, respectively. In many instances, having a high stiffness (Elastic Modulus) can predict the ability to withstand higher stresses. There was also a low correlation of Elastic Modulus and Peak Stress to ACTA2 (0.33 and 0.32, respectfully) and TGFβ1 (0.28 and 0.27, respectfully), which are associated with myofibroblast activation and collagen secretion [120]. Elastic modulus was also a key contributor to tissue variability and contributed the most to the fifth dimension in the FAMD analysis (**Table 2**), demonstrating that tissue mechanics strongly affects other tissue properties.

Both the Elastic Modulus and Peak Stress variables had a low correlation with hydroxyproline content (0.35 and 0.30, respectfully) which follows similar trends, with several exceptions. In general, increased hydroxyproline content, the main component of collagen, was paired with increased elastic moduli and peak stress, with the exception of patient 13 and 15. One reason for these differences between mechanical properties and collagen content is the lack of specificity in the assay. Hydroxyproline assay kits do not differentiate between different types of collagen but simply measure the total hydroxyproline content. Collagens, such as type I, III, IV, V, and VI, are found in adipose tissue and have been linked to obesity and adipose tissue fibrosis [26, 121–123]. However, the role of these collagens varies. Type IV and VI are network-forming collagens found in the basement membrane, while type I, III, and V are fibril-forming collagens found in the interstitial space [124]. Fibrosis can affect the relative concentration of interstitial collagens to basement membrane collagens changing the mechanical properties [125].

To evaluate cellular metabolism, we used resazurin which is transformed from a blue color to the resorufin (pink) by a redox reaction process. Patient 10 had the smallest average adipocyte diameter and the highest metabolic activity while patient 12 had the largest average adipocyte diameter and had a low redox indicator. However, it should be noted that the relationship between adipocyte diameter and what we are defining as “metabolic activity” was not strongly correlated in our data (−0.22). One study indicated that small adipocytes are insufficient to protect against metabolic dysfunction [126] and that any increase in adipocyte size, either from small or normal size cells, could result in reduced metabolism [3, 126]. However, there was a positive correlation between metabolic activity and advanced glycation end products (0.48) supporting literature that shows advanced glycation end products regulate extracellular matrix-adipocyte metabolic crosstalk [127]. Metabolic activity was also negatively correlated with CD86 (M1 macrophage marker, −0.41) indicating adipose tissue metabolism decreased with a proinflammatory M1 macrophage phenotype.

Collectively looking at this dataset, Patient 15 and 16 offer a unique comparison. Both patients are female, age 48, diabetic, non-smoking, with similar BMIs. The only difference between them is that patient 16 has a history of obesity. They do not have statistically different elastic moduli or peak stress but patient 16 has a significantly quicker doubling time and higher metabolic activity. Patient 15 also has a higher hydroxyproline content.

**Supplemental Figure 3.**
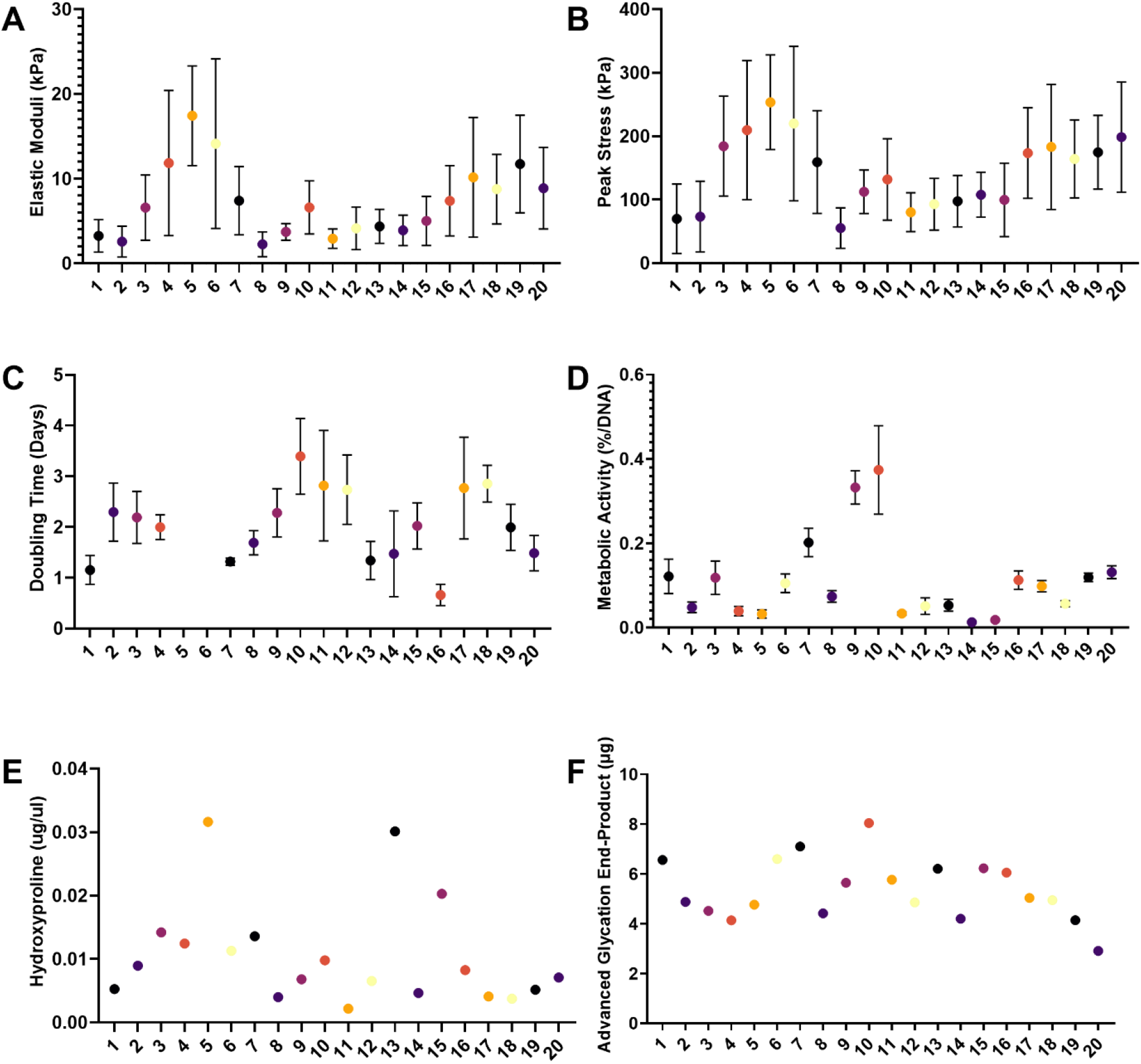
(A) Elastic moduli, (B) Peak Stress, (C) SVF Doubling Time, (D) Metabolic Activity, (E) Hydroxyproline, and (F) Advanced glycation end-product concentration for each patient in the study.

**Supplemental Figure 4.**
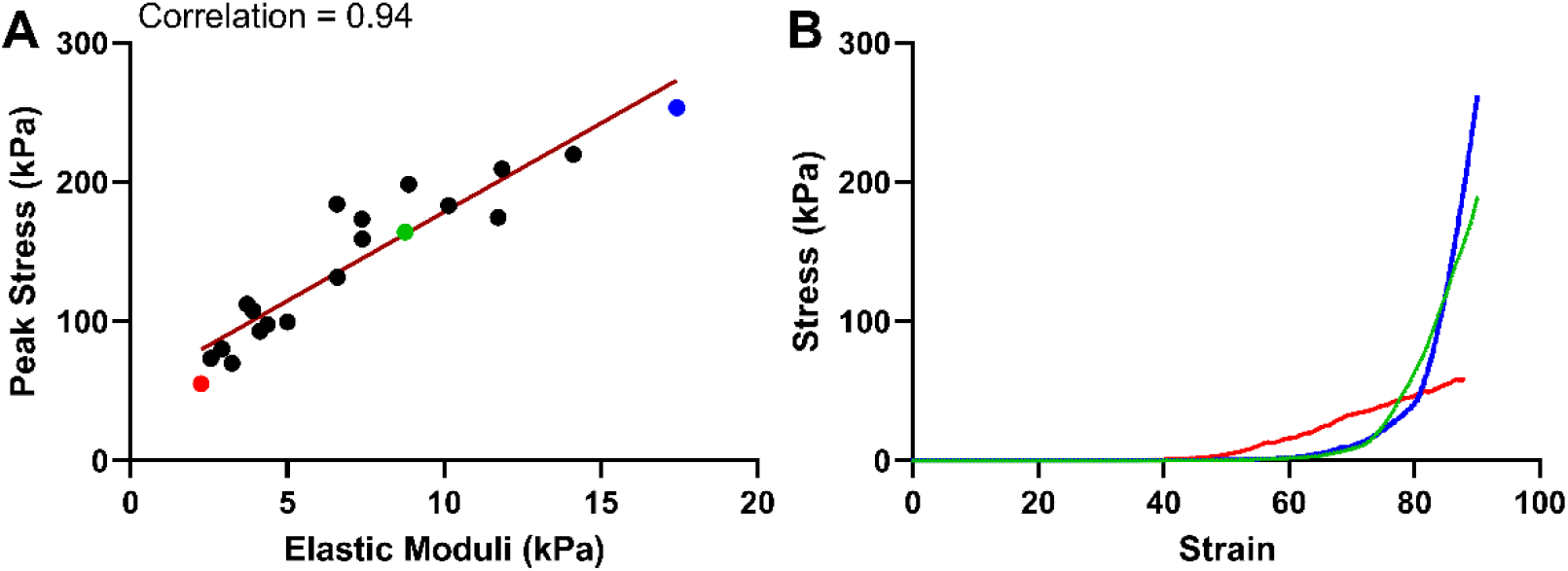
Elastic modulus and peak stress are highly correlated. The highest correlation in the dataset was between Elastic Modulus and Peak Stress (0.94). Peak stress was plotted versus Elastic Modulus for the 20 patients (A), where 3 example plots of stress versus strain are shown (B) that correspond to the same colors highlighted in A (red, green, and blue).

**Supplemental Table 3.** Statistical significance for elastic modulus. Determined through a one-way ANOVA followed by Tukey’s post-hoc analysis. Non-statistically significant comparisons are not shown. Significance was defined as p<0.05. * p<0.05, ** p<0.01, *** p<0.005, **** p<0.0001.

| Comparison | Significance | P value | Comparison | Significance | P value |
| --- | --- | --- | --- | --- | --- |
| 1 vs. 9 | * | 0.025 | 6 vs. 11 | * | 0.0105 |
| 1 vs. 11 | ** | 0.0017 | 6 vs. 17 | * | 0.0232 |
| 1 vs. 12 | * | 0.0399 | 6 vs. 18 | **** | <0.0001 |
| 1 vs. 16 | * | 0.0251 | 6 vs. 19 | **** | <0.0001 |
| 1 vs. 17 | ** | 0.0042 | 7 vs. 18 | * | 0.0466 |
| 1 vs. 18 | **** | <0.0001 | 7 vs. 19 | * | 0.02 |
| 1 vs. 19 | **** | <0.0001 | 8 vs. 9 | * | 0.0274 |
| 2 vs. 4 | * | 0.0397 | 8 vs. 11 | ** | 0.0019 |
| 2 vs. 9 | ** | 0.0011 | 8 vs. 12 | * | 0.0436 |
| 2 vs. 11 | **** | <0.0001 | 8 vs. 16 | * | 0.0275 |
| 2 vs. 12 | ** | 0.002 | 8 vs. 17 | ** | 0.0047 |
| 2 vs. 15 | * | 0.0112 | 8 vs. 18 | **** | <0.0001 |
| 2 vs. 16 | ** | 0.0011 | 8 vs. 19 | **** | <0.0001 |
| 2 vs. 17 | *** | 0.0001 | 10 vs. 18 | * | 0.0349 |
| 2 vs. 18 | **** | <0.0001 | 10 vs. 19 | * | 0.0145 |
| 2 vs. 19 | **** | <0.0001 | 11 vs. 14 | ** | 0.0053 |
| 2 vs. 20 | ** | 0.0038 | 13 vs. 18 | ** | 0.0057 |
| 3 vs. 19 | * | 0.026 | 13 vs. 19 | ** | 0.002 |
| 5 vs. 18 | ** | 0.0029 | 14 vs. 17 | * | 0.0122 |
| 5 vs. 19 | *** | 0.001 | 14 vs. 18 | **** | <0.0001 |
|  |  |  | 14 vs. 19 | **** | <0.0001 |

**Supplemental Table 4.**
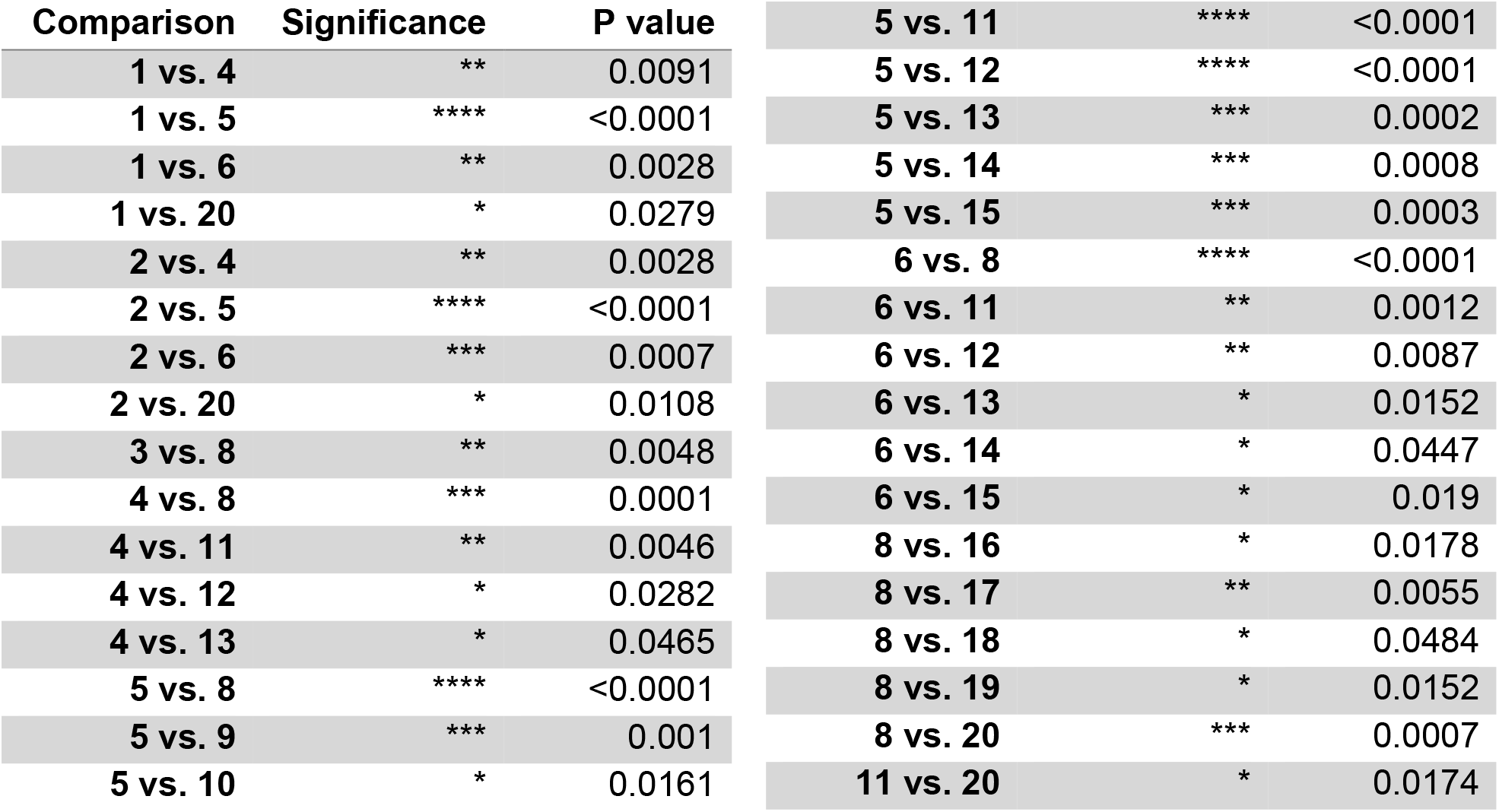
Statistical significance for peak stress. Determined through a one-way ANOVA followed by Tukey’s post-hoc analysis. Non-statistically significant comparisons are not shown. Significance was defined as p<0.05. * p<0.05, ** p<0.01, *** p<0.005, **** p<0.0001.

**Supplemental Table 5.** Statistical significance for SVF doubling time. Determined through a one-way ANOVA followed by Tukey’s post-hoc analysis. Non-statistically significant comparisons are not shown. Significance was defined as p<0.05. * p<0.05, ** p<0.01, *** p<0.005, **** p<0.0001.

| Comparison | Significance | P value |
| --- | --- | --- |
| 1 vs. 10 | **** | <0.0001 |
| 1 vs. 11 | ** | 0.0019 |
| 1 vs. 12 | ** | 0.0042 |
| 1 vs. 17 | ** | 0.0031 |
| 1 vs. 18 | ** | 0.0013 |
| 2 vs. 16 | ** | 0.0026 |
| 3 vs. 16 | ** | 0.0069 |
| 4 vs. 10 | * | 0.0226 |
| 4 vs. 16 | * | 0.0369 |
| 7 vs. 10 | **** | <0.0001 |
| 7 vs. 11 | ** | 0.0093 |
| 7 vs. 12 | * | 0.0188 |
| 7 vs. 17 | * | 0.0142 |
| 7 vs. 18 | ** | 0.0066 |
| 8 vs. 10 | ** | 0.0013 |
| 9 vs. 16 | ** | 0.0029 |
| 10 vs. 13 | **** | <0.0001 |
| 10 vs. 14 | *** | 0.0001 |
| 10 vs. 15 | * | 0.0284 |
| 10 vs. 16 | **** | <0.0001 |
| 10 vs. 19 | * | 0.0221 |
| 10 vs. 20 | *** | 0.0002 |
| 11 vs. 13 | * | 0.0112 |
| 11 vs. 14 | * | 0.0348 |
| 11 vs. 16 | **** | <0.0001 |
| 11 vs. 20 | * | 0.0388 |
| 12 vs. 13 | * | 0.0226 |
| 12 vs. 16 | **** | <0.0001 |
| 13 vs. 17 | * | 0.017 |
| 13 vs. 18 | ** | 0.008 |
| 14 vs. 18 | * | 0.0255 |
| 15 vs. 16 | * | 0.0296 |
| 16 vs. 17 | **** | <0.0001 |
| 16 vs. 18 | **** | <0.0001 |
| 16 vs. 19 | * | 0.0377 |
| 18 vs. 20 | * | 0.0286 |

**Supplemental Table 6.**
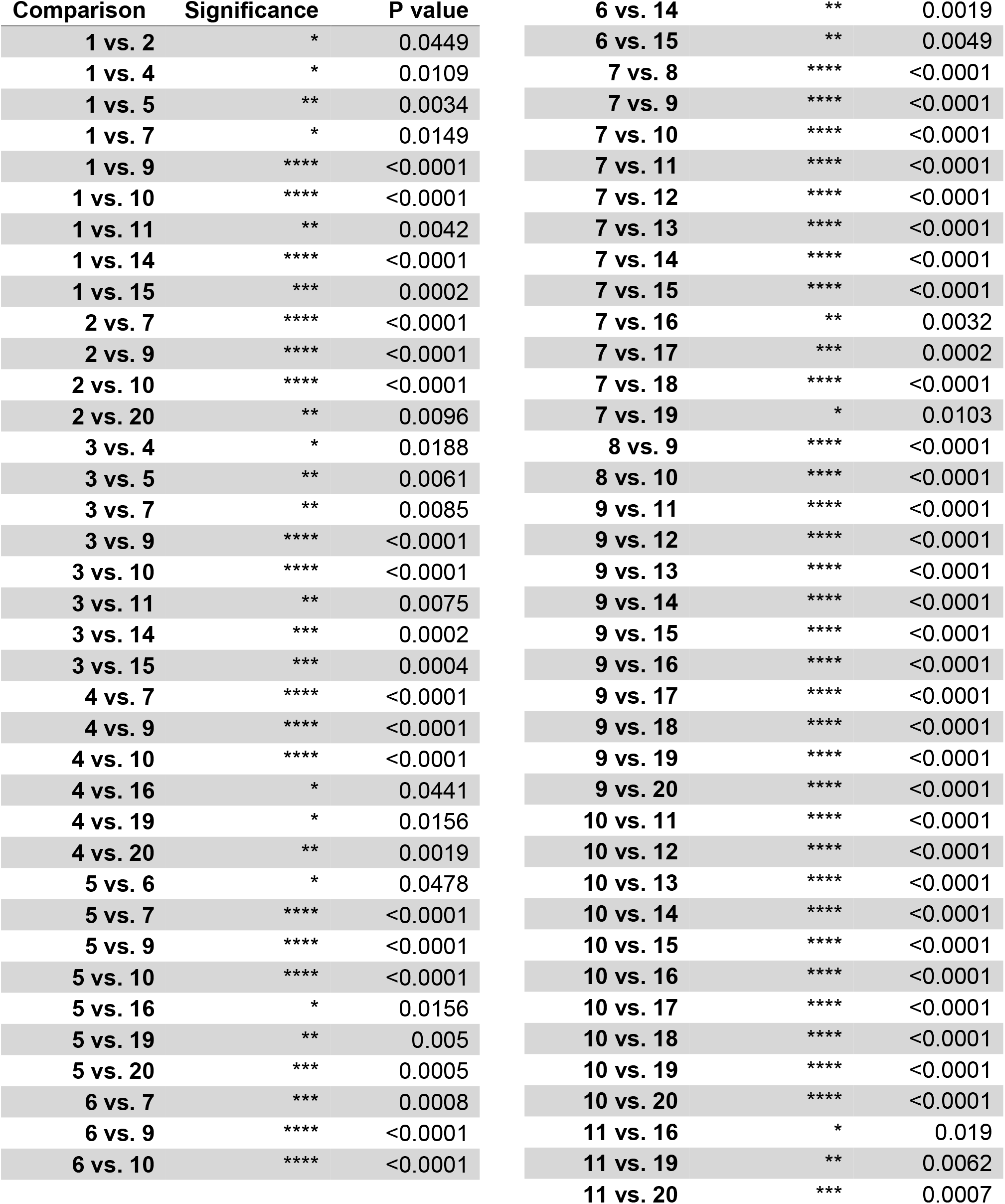
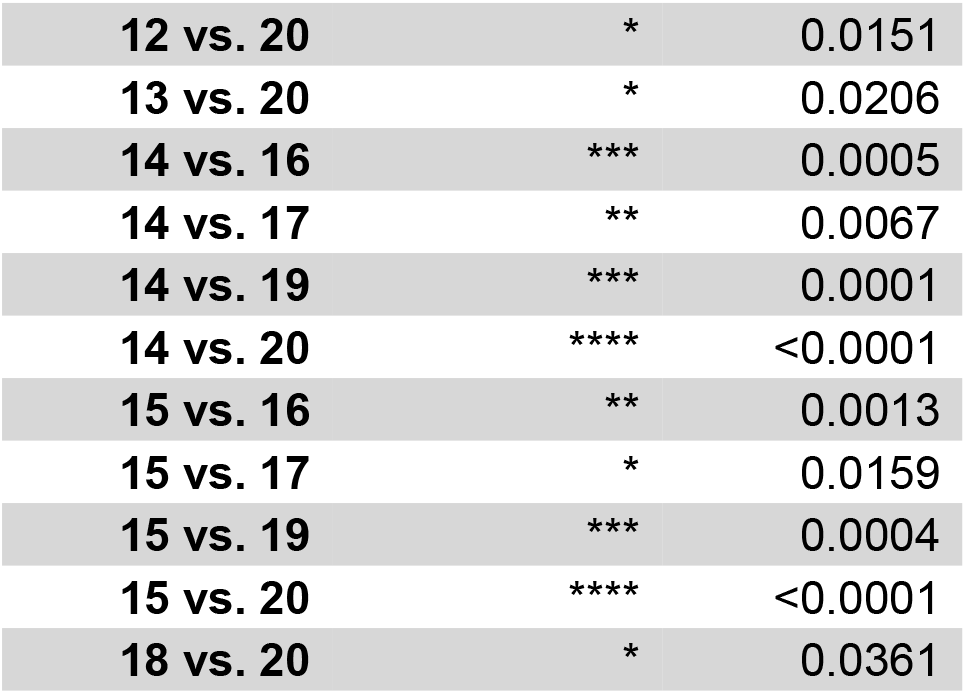
Statistical significance for metabolic activity (resazurin). Determined through a one-way ANOVA followed by Tukey’s post-hoc analysis. Non-statistically significant comparisons are not shown. Significance was defined as p<0.05. * p<0.05, ** p<0.01, *** p<0.005, **** p<0.0001.

| Comparison | Significance | P value |  |  |  |
| --- | --- | --- | --- | --- | --- |
| 1 vs. 2 | * | 0.0449 | 6 vs. 14 | ** | 0.0019 |
| 1 vs. 4 | * | 0.0109 | 6 vs. 15 | ** | 0.0049 |
| 1 vs. 5 | ** | 0.0034 | 7 vs. 8 | **** | <0.0001 |
| 1 vs. 7 | * | 0.0149 | 7 vs. 9 | **** | <0.0001 |
| 1 vs. 9 | **** | <0.0001 | 7 vs. 10 | **** | <0.0001 |
| 1 vs. 10 | **** | <0.0001 | 7 vs. 11 | **** | <0.0001 |
| 1 vs. 11 | ** | 0.0042 | 7 vs. 12 | **** | <0.0001 |
| 1 vs. 14 | **** | <0.0001 | 7 vs. 13 | **** | <0.0001 |
| 1 vs. 15 | *** | 0.0002 | 7 vs. 14 | **** | <0.0001 |
| 2 vs. 7 | **** | <0.0001 | 7 vs. 15 | **** | <0.0001 |
| 2 vs. 9 | **** | <0.0001 | 7 vs. 16 | ** | 0.0032 |
| 2 vs. 10 | **** | <0.0001 | 7 vs. 17 | *** | 0.0002 |
| 2 vs. 20 | ** | 0.0096 | 7 vs. 18 | **** | <0.0001 |
| 3 vs. 4 | * | 0.0188 | 7 vs. 19 | * | 0.0103 |
| 3 vs. 5 | ** | 0.0061 | 8 vs. 9 | **** | <0.0001 |
| 3 vs. 7 | ** | 0.0085 | 8 vs. 10 | **** | <0.0001 |
| 3 vs. 9 | **** | <0.0001 | 9 vs. 11 | **** | <0.0001 |
| 3 vs. 10 | **** | <0.0001 | 9 vs. 12 | **** | <0.0001 |
| 3 vs. 11 | ** | 0.0075 | 9 vs. 13 | **** | <0.0001 |
| 3 vs. 14 | *** | 0.0002 | 9 vs. 14 | **** | <0.0001 |
| 3 vs. 15 | *** | 0.0004 | 9 vs. 15 | **** | <0.0001 |
| 4 vs. 7 | **** | <0.0001 | 9 vs. 16 | **** | <0.0001 |
| 4 vs. 9 | **** | <0.0001 | 9 vs. 17 | **** | <0.0001 |
| 4 vs. 10 | **** | <0.0001 | 9 vs. 18 | **** | <0.0001 |
| 4 vs. 16 | * | 0.0441 | 9 vs. 19 | **** | <0.0001 |
| 4 vs. 19 | * | 0.0156 | 9 vs. 20 | **** | <0.0001 |
| 4 vs. 20 | ** | 0.0019 | 10 vs. 11 | **** | <0.0001 |
| 5 vs. 6 | * | 0.0478 | 10 vs. 12 | **** | <0.0001 |
| 5 vs. 7 | **** | <0.0001 | 10 vs. 13 | **** | <0.0001 |
| 5 vs. 9 | **** | <0.0001 | 10 vs. 14 | **** | <0.0001 |
| 5 vs. 10 | **** | <0.0001 | 10 vs. 15 | **** | <0.0001 |
| 5 vs. 16 | * | 0.0156 | 10 vs. 16 | **** | <0.0001 |
| 5 vs. 19 | ** | 0.005 | 10 vs. 17 | **** | <0.0001 |
| 5 vs. 20 | *** | 0.0005 | 10 vs. 18 | **** | <0.0001 |
| 6 vs. 7 | *** | 0.0008 | 10 vs. 19 | **** | <0.0001 |
| 6 vs. 9 | **** | <0.0001 | 10 vs. 20 | **** | <0.0001 |
| 6 vs. 10 | **** | <0.0001 | 11 vs. 16 | * | 0.019 |
|  |  |  | 11 vs. 19 | ** | 0.0062 |
|  |  |  | 11 vs. 20 | *** | 0.0007 |

|  |  |  |
| --- | --- | --- |
| **12 vs. 20** | **\*** | **0.0151** |
| **13 vs. 20** | **\*** | **0.0206** |
| **14 vs. 16** | **\*\*\*** | **0.0005** |
| **14 vs. 17** | **\*\*** | **0.0067** |
| **14 vs. 19** | **\*\*\*** | **0.0001** |
| **14 vs. 20** | **\*\*\*\*** | **<0.0001** |
| **15 vs. 16** | **\*\*** | **0.0013** |
| **15 vs. 17** | **\*** | **0.0159** |
| **15 vs. 19** | **\*\*\*** | **0.0004** |
| **15 vs. 20** | **\*\*\*\*** | **<0.0001** |
| **18 vs. 20** | **\*** | **0.0361** |

### Supplement 3. Gene Expression

SDHA was used as our reference gene due to its high stability in human subcutaneous adipose tissue [31]. Because no control group was used in this study gene expression is expressed as ΔCT values:

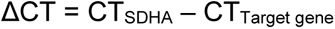

Using this formula higher CT values of the target gene, which indicate lower gene expression, would have lower ΔCT values.

Both IL6 and TNFα exhibit higher gene expression in metabolically dysfunctional adipose tissue [26] and were correlated in this dataset (correlation = 0.45). For most patients, a higher IL6 gene expression (**Supplemental Figure 5A**) correlated with a higher TNFα gene expression (**Supplemental Figure 5B**). Interestingly patient 8, the only patient to have an IL6 CT value over 40, also had the lowest TNFα gene expression. TNFα is also linked to increased intracellular lipid accumulation [128]. Interestingly, patients 2, 13, and 14 exhibited increased expression of TNFα expression and also an increased number of extracellular lipid droplets (**Supplemental Figure 1**).

TGFβ1 expression is increased during obesity [129] and when expression is blocked can prevent obesity, insulin resistance, and fatty liver disease [130]. As expected, the patients exhibiting the highest TGFβ1 expression (**Supplemental Figure 5E**) currently have diabetes or are pre-diabetic, which were patients 9, 13 and 16. Older patients, like patients 2, 3, and 17, also experienced an increase in TGFβ1 expression.

Adiponectin inhibits inflammation, which has been shown to reduce collagen content in murine adipose tissue [25, 131]. It is exclusively produced by adipocytes. Adiponectin expression is lower in obese and diabetic individuals and increased during weight loss [132]. Due to the nature of the procedure, it is known that the patients that underwent a panniculectomy procedure lost a significant amount of weight, but the time from weight loss is not known. The individuals that have diabetes have a range of gene expression patterns with patients 1 and 8 having decreased gene expression compared to patients 9, 15, 16, and 20 that all experienced increased gene expression (**Supplemental Figure 5I**). Additionally, patient 20 has the highest BMI of the patients we studied.

Leptin has been shown to increase collagen type I content and stimulate TGFβ1 expression [133]. Our results do indicate that patients with higher leptin levels (**Supplemental Figure 5D)** also experienced higher TGFβ1 gene expression (**Supplemental Figure 5C**). However, the leptin levels do not correlate with higher collagen levels. Patients 5 and 13 had the highest hydroxyproline content (**Supplemental Figure 3E**) but have lower leptin gene expression than the majority of other patients.

**Supplemental Figure 5.**
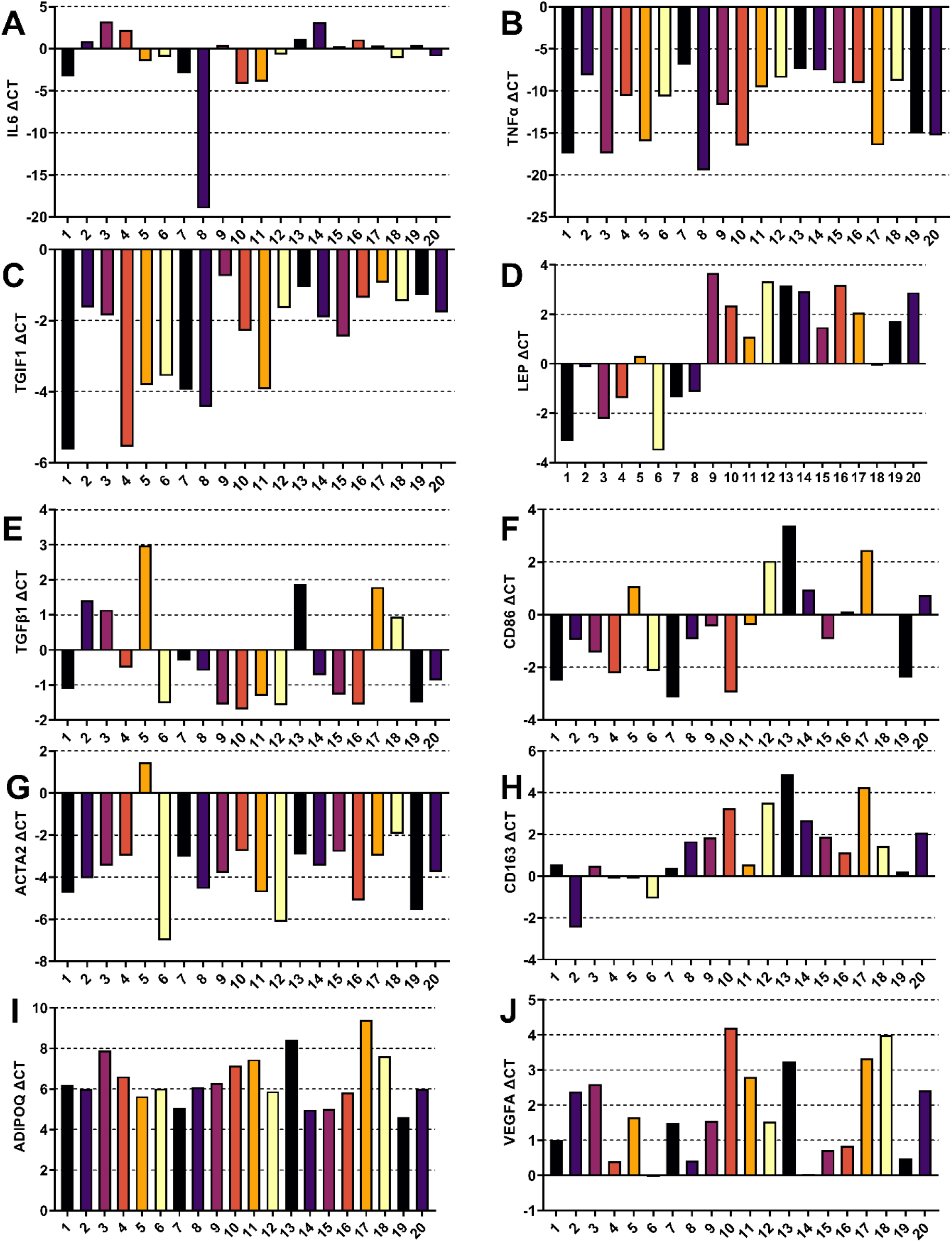
Gene expression for each patient shown by the ΔCT value.

### Supplement 4. FAMD coordinates

**Supplemental Figure 6.**
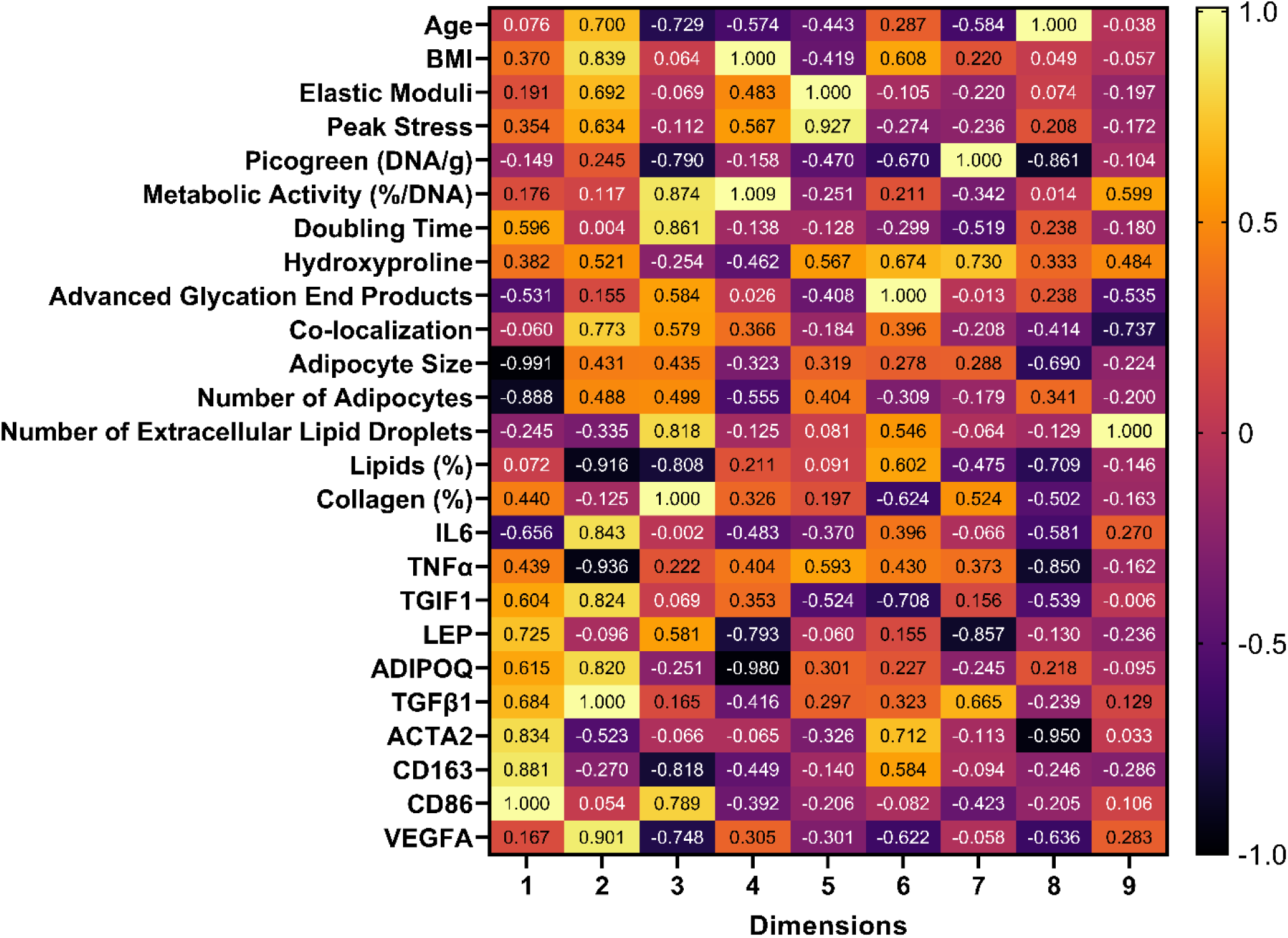
Quantitative variable coordinates in each dimension generated from the FAMD analysis normalized to the furthest coordinate in each dimension. High positive coordinate values indicate the variables are clustered and more similar (for example in the first dimension ACTA2, CD86, and CD163), while a negative value would indicate the variables are dissimilar and ordinated further away from that cluster (for example Adipocyte Size and Number of Adipocytes).

**Supplement Figure 7.**
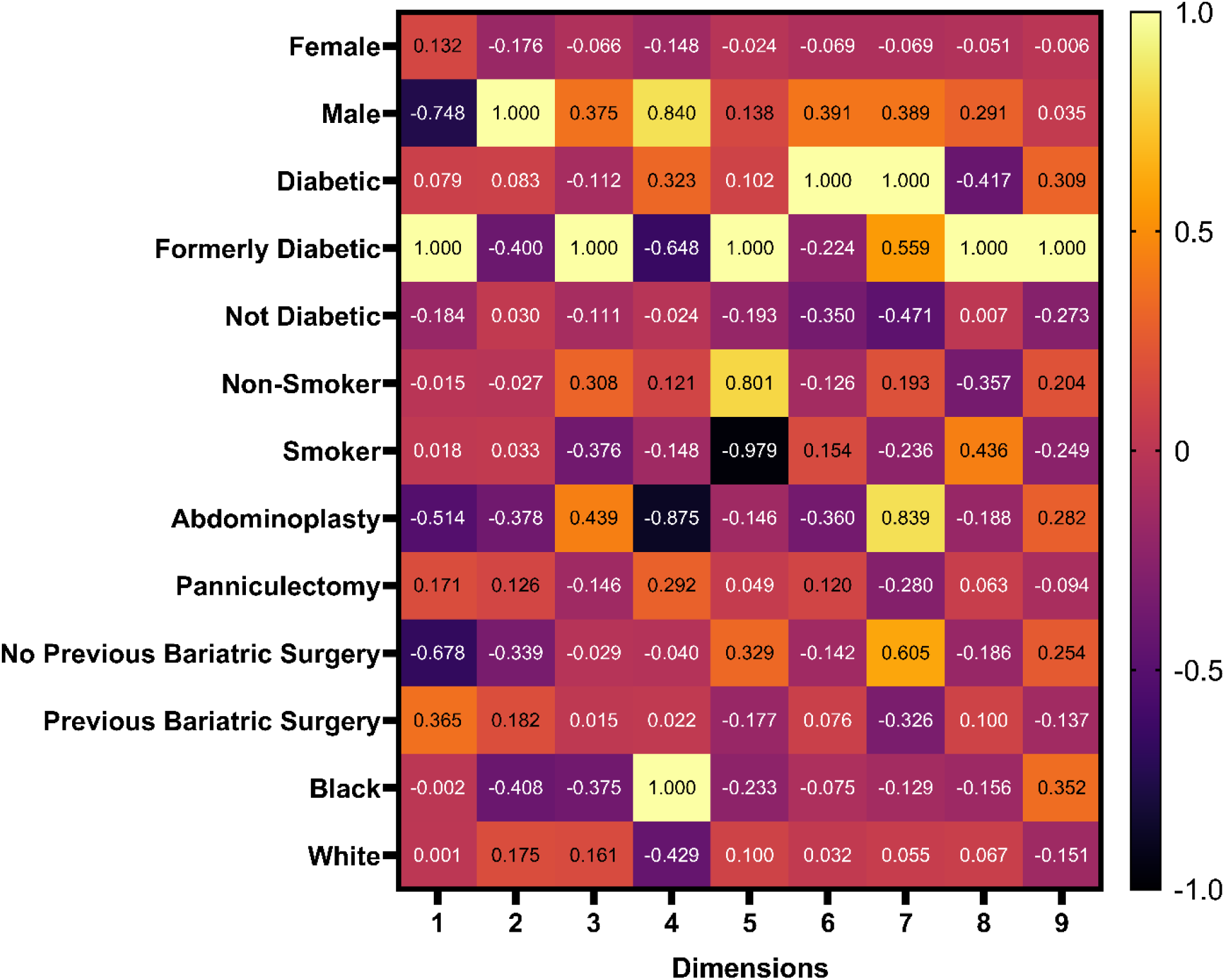
Qualitative variable coordinates generated from the FAMD analysis normalized to the furthest coordinate in each dimension.

### Supplement 5. Code used for running FAMD

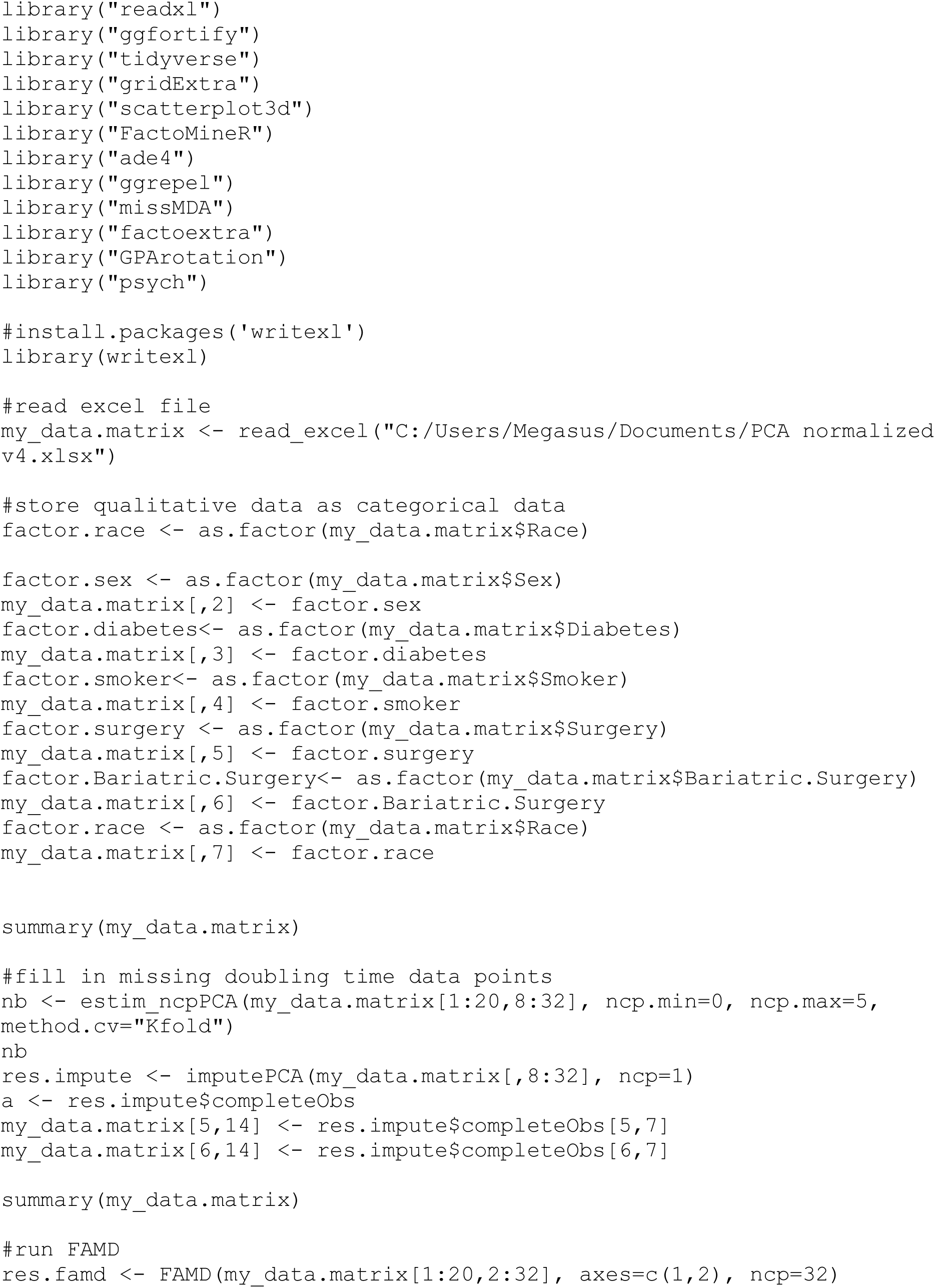
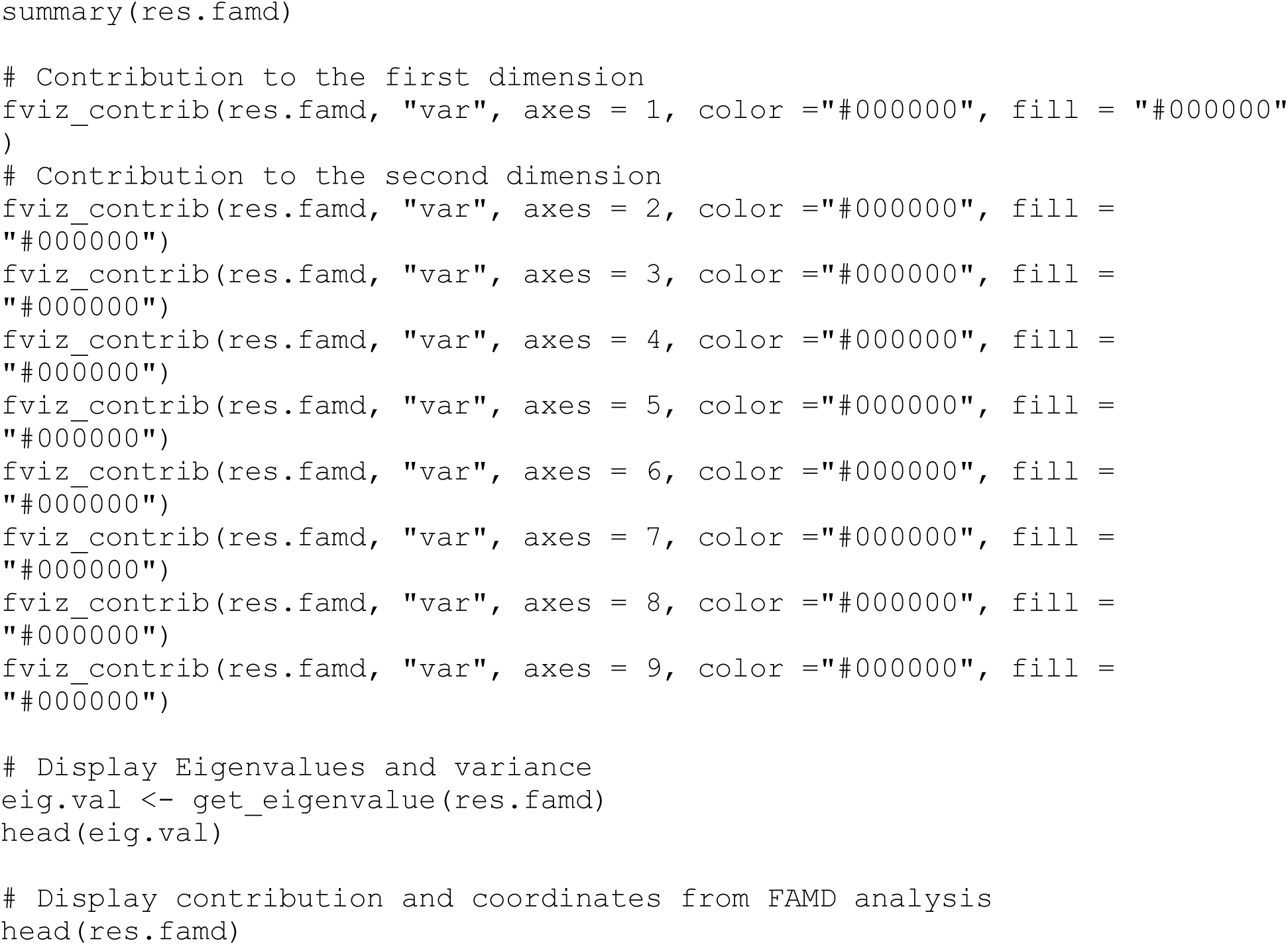

## References

1. Wolin, K. Y.; Carson, K.; Colditz, G. A., Obesity and cancer. The oncologist 2010, 15 (6), 556–65.

2. Barnes, A. S., The epidemic of obesity and diabetes: trends and treatments. Texas Heart Institute journal 2011, 38 (2), 142–4.

3. Stenkula, K. G.; Erlanson-Albertsson, C., Adipose cell size: importance in health and disease. *American Journal of Physiology-Regulatory*, Integrative and Comparative Physiology 2018, 315 (2), R284–R295.

4. Abbott, R. D.; Borowsky, F. E.; Alonzo, C. A.; Zieba, A.; Georgakoudi, I.; Kaplan, D. L., Variability in responses observed in human white adipose tissue models. J Tissue Eng Regen Med 2018, 12 (3), 840–847.

5. Alkhouli, N.; Mansfield, J.; Green, E.; Bell, J.; Knight, B.; Liversedge, N.; Tham, J. C.; Welbourn, R.; Shore, A. C.; Kos, K.; Winlove, C. P., The mechanical properties of human adipose tissues and their relationships to the structure and composition of the extracellular matrix. Am J Physiol Endocrinol Metab 2013, 305 (12), E1427–35.

6. Divoux, A.; Tordjman, J.; Lacasa, D.; Veyrie, N.; Hugol, D.; Aissat, A.; Basdevant, A.; Guerre-Millo, M.; Poitou, C.; Zucker, J.-D.; Bedossa, P.; Clément, K., Fibrosis in Human Adipose Tissue: Composition, Distribution, and Link With Lipid Metabolism and Fat Mass Loss. Diabetes 2010, 59 (11), 2817–2825.

7. Yi, C. G.; Pan, Y.; Zhen, Y.; Zhang, L. X.; Zhang, X. D.; Shu, M. G.; Han, Y.; Guo, S. Z., Enhancement of viability of fat grafts in nude mice by endothelial progenitor cells. Dermatol Surg 2006, 32 (12), 1437–1443.

8. Rieck, B.; Schlaak, S., Measurement in vivo of the survival rate in autologous adipocyte transplantation. Plast Reconstr Surg 2003, 111 (7), 2315–2323.

9. Meier, J. D.; Glasgold, R. A.; Glasgold, M. J., Autologous Fat Grafting Long-term Evidence of Its Efficacy in Midfacial Rejuvenation. Arch Facial Plast S 2009, 11 (1), 24–28.

10. Report on autologous fat transplantation. ASPRS Ad-Hoc Committee on New Procedures, September 30, 1987. Plast Surg Nurs 1987, 7 (4), 140–1.

11. Chabot, K.; Gauthier, M. S.; Garneau, P. Y.; Rabasa-Lhoret, R., Evolution of subcutaneous adipose tissue fibrosis after bariatric surgery. Diabetes Metab 2017, 43 (2), 125–133.

12. Muir, L. A.; Neeley, C. K.; Meyer, K. A.; Baker, N. A.; Brosius, A. M.; Washabaugh, A. R.; Varban, O. A.; Finks, J. F.; Zamarron, B. F.; Flesher, C. G.; Chang, J. S.; DelProposto, J. B.; Geletka, L.; Martinez-Santibanez, G.; Kaciroti, N.; Lumeng, C. N.; O’Rourke, R. W., Adipose tissue fibrosis, hypertrophy, and hyperplasia: Correlations with diabetes in human obesity. Obesity (Silver Spring) 2016, 24 (3), 597–605.

13. Anand, S. S.; Tarnopolsky, M. A.; Rashid, S.; Schulze, K. M.; Desai, D.; Mente, A.; Rao, S.; Yusuf, S.; Gerstein, H. C.; Sharma, A. M., Adipocyte hypertrophy, fatty liver and metabolic risk factors in South Asians: the Molecular Study of Health and Risk in Ethnic Groups (mol-SHARE). PLoS One 2011, 6 (7), e22112.

14. Gu, D.; He, J.; Duan, X.; Reynolds, K.; Wu, X.; Chen, J.; Huang, G.; Chen, C. S.; Whelton, P. K., Body weight and mortality among men and women in China. JAMA 2006, 295 (7), 776–83.

15. Alba, D. L.; Farooq, J. A.; Lin, M. Y. C.; Schafer, A. L.; Shepherd, J.; Koliwad, S. K., Subcutaneous Fat Fibrosis Links Obesity to Insulin Resistance in Chinese Americans. J Clin Endocrinol Metab 2018, 103 (9), 3194–3204.

16. Jan, V.; Cervera, P.; Maachi, M.; Baudrimont, M.; Kim, M.; Vidal, H.; Girard, P. M.; Levan, P.; Rozenbaum, W.; Lombès, A.; Capeau, J.; Bastard, J. P., Altered fat differentiation and adipocytokine expression are inter-related and linked to morphological changes and insulin resistance in HIV-1-infected lipodystrophic patients. Antiviral therapy 2004, 9 (4), 555–64.

17. Han, S.; Sun, H. M.; Hwang, K. C.; Kim, S. W., Adipose-Derived Stromal Vascular Fraction Cells: Update on Clinical Utility and Efficacy. Crit Rev Eukaryot Gene Expr 2015, 25 (2), 145–52.

18. Jin Young, H.; Yoon Jeong, P.; Mira, H.; and Jae Bum, K.; Crosstalk between Adipocytes and Immune Cells in Adipose Tissue Inflammation and Metabolic Dysregulation in Obesity. Mol. Cells 2014, 37 (5), 365–371.

19. Verboven, K.; Wouters, K.; Gaens, K.; Hansen, D.; Bijnen, M.; Wetzels, S.; Stehouwer, C. D.; Goossens, G. H.; Schalkwijk, C. G.; Blaak, E. E.; Jocken, J. W., Abdominal subcutaneous and visceral adipocyte size, lipolysis and inflammation relate to insulin resistance in male obese humans. Sci Rep 2018, 8 (1), 4677.

20. Drolet, R.; Richard, C.; Sniderman, A. D.; Mailloux, J.; Fortier, M.; Huot, C.; Rhéaume, C.; Tchernof, A., Hypertrophy and hyperplasia of abdominal adipose tissues in women. International journal of obesity (2005) 2008, 32 (2), 283–91.

21. Von Bank, H.; Kirsh, C.; Simcox, J., Aging adipose: Depot location dictates age-associated expansion and dysfunction. Ageing research reviews 2021, 67, 101259.

22. Muir, L. A.; Neeley, C. K.; Meyer, K. A.; Baker, N. A.; Brosius, A. M.; Washabaugh, A. R.; Varban, O. A.; Finks, J. F.; Zamarron, B. F.; Flesher, C. G., Adipose tissue fibrosis, hypertrophy, and hyperplasia: Correlations with diabetes in human obesity. Obesity 2016, 24 (3), 597–605.

23. Arner, E.; Westermark, P. O.; Spalding, K. L.; Britton, T.; Rydén, M.; Frisén, J.; Bernard, S.; Arner, P., Adipocyte turnover: relevance to human adipose tissue morphology. Diabetes 2010, 59 (1), 105–109.

24. Ali, A. T.; Hochfeld, W. E.; Myburgh, R.; Pepper, M. S., Adipocyte and adipogenesis. European journal of cell biology 2013, 92 (6-7), 229–236.

25. Khan, T.; Muise, E. S.; Iyengar, P.; Wang, Z. V.; Chandalia, M.; Abate, N.; Zhang, B. B.; Bonaldo, P.; Chua, S.; Scherer, P. E., Metabolic dysregulation and adipose tissue fibrosis: role of collagen VI. Molecular and cellular biology 2009, 29 (6), 1575–91.

26. DeBari, M. K.; Abbott, R. D., Adipose Tissue Fibrosis: Mechanisms, Models, and Importance. Int J Mol Sci 2020, 21 (17).

27. Shook, B.; Rodeheffer, M. S., Forecasting Fat Fibrosis. Cell Metabolism 2017, 25 (3), 493–494.

28. Crewe, C.; An, Y. A.; Scherer, P. E., The ominous triad of adipose tissue dysfunction: inflammation, fibrosis, and impaired angiogenesis. The Journal of clinical investigation 2017, 127 (1), 74–82.

29. Comley, K.; Fleck, N., The compressive response of porcine adipose tissue from low to high strain rate. International Journal of Impact Engineering 2012, 46, 1–10.

30. Abbott, R. D.; Raja, W. K.; Wang, R. Y.; Stinson, J. A.; Glettig, D. L.; Burke, K. A.; Kaplan, D. L., Long term perfusion system supporting adipogenesis. Methods 2015, 84, 84–9.

31. Perez, L. J.; Rios, L.; Trivedi, P.; D’Souza, K.; Cowie, A.; Nzirorera, C.; Webster, D.; Brunt, K.; Legare, J. F.; Hassan, A.; Kienesberger, P. C.; Pulinilkunnil, T., Validation of optimal reference genes for quantitative real time PCR in muscle and adipose tissue for obesity and diabetes research. Sci Rep 2017, 7 (1), 3612.

32. Carre, A.; Klausner, G.; Edjlali, M.; Lerousseau, M.; Briend-Diop, J.; Sun, R.; Ammari, S.; Reuze, S.; Alvarez Andres, E.; Estienne, T.; Niyoteka, S.; Battistella, E.; Vakalopoulou, M.; Dhermain, F.; Paragios, N.; Deutsch, E.; Oppenheim, C.; Pallud, J.; Robert, C., Standardization of brain MR images across machines and protocols: bridging the gap for MRI-based radiomics. Sci Rep 2020, 10 (1), 12340.

33. Cheadle, C.; Vawter, M. P.; Freed, W. J.; Becker, K. G., Analysis of microarray data using Z score transformation. J Mol Diagn 2003, 5 (2), 73–81.

34. Josse, J.; Husson, F., missMDA: A Package for Handling Missing Values in Multivariate Data Analysis. Journal of Statistical Software 2016, 70 (1).

35. Johnston, E. K.; Abbott, R. D., Adipose Tissue Paracrine-, Autocrine-, and Matrix-Dependent Signaling during the Development and Progression of Obesity. Cells 2023, 12 (3), 407.

36. Johnston, E. K.; Abbott, R. D., Adipose Tissue Development Relies on Coordinated Extracellular Matrix Remodeling, Angiogenesis, and Adipogenesis. Biomedicines 2022, 10 (9), 2227.

37. Elias, I.; Franckhauser, S.; Bosch, F., New insights into adipose tissue VEGF-A actions in the control of obesity and insulin resistance. Adipocyte 2013, 2 (2), 109–12.

38. Sun, K.; Asterholm, I. W.; Kusminski, C. M.; Bueno, A. C.; Wang, Z. V.; Pollard, J. W.; Brekken, R. A.; Scherer, P. E., Dichotomous effects of VEGF-A on adipose tissue dysfunction. Proceedings of the national academy of sciences 2012, 109 (15), 5874–5879.

39. Lemoine, A. Y.; Ledoux, S.; Larger, E., Adipose tissue angiogenesis in obesity. Thrombosis and haemostasis 2013, 110 (10), 661–669.

40. Zuk, P. A.; Zhu, M.; Mizuno, H.; Huang, J.; Futrell, J. W.; Katz, A. J.; Benhaim, P.; Lorenz, H. P.; Hedrick, M. H., Multilineage cells from human adipose tissue: implications for cell-based therapies. Tissue engineering 2001, 7 (2), 211–228.

41. Moegni, K.; Rosliana, I.; Sobariah, S.; Rosadi, I.; Afini, I.; Widyastuti, T.; Remelia, M.; Sukmawati, D.; Pawitan, J., Diabetes mellitus type 2 reduces the viability, proliferation, and angiogenic marker of adipose-derived stem cells cultured in low-glucose anti-oxidant-serum supplemented medium. Biomedical Research and Therapy 2019, 6, 3073–3082.

42. Yoshimura, K.; Shigeura, T.; Matsumoto, D.; Sato, T.; Takaki, Y.; Aiba-Kojima, E.; Sato, K.; Inoue, K.; Nagase, T.; Koshima, I., Characterization of freshly isolated and cultured cells derived from the fatty and fluid portions of liposuction aspirates. J Cell Physiol 2006, 208 (1), 64–76.

43. Alt, E. U.; Senst, C.; Murthy, S. N.; Slakey, D. P.; Dupin, C. L.; Chaffin, A. E.; Kadowitz, P. J.; Izadpanah, R., Aging alters tissue resident mesenchymal stem cell properties. Stem cell research 2012, 8 (2), 215–225.

44. Perez, L. M.; Bernal, A.; de Lucas, B.; San Martin, N.; Mastrangelo, A.; Garcia, A.; Barbas, C.; Galvez, B. G., Altered metabolic and stemness capacity of adipose tissue-derived stem cells from obese mouse and human. PLoS One 2015, 10 (4), e0123397.

45. Ruhrberg, C.; Gerhardt, H., VEGF and endothelial guidance in angiogenic sprouting. VEGF in Development 2008, 68–78.

46. Fukumura, D.; Ushiyama, A.; Duda, D. G.; Xu, L.; Tam, J.; Chatterjee, V. K. K.; Garkavtsev, I.; Jain, R. K., Paracrine regulation of angiogenesis and adipocyte differentiation during in vivo adipogenesis. Circ Res 2003, 93 (9), E88–E97.

47. Fukumura, D.; Ushiyama, A.; Duda, D. G.; Xu, L.; Tam, J.; Krishna, V.; Chatterjee, K.; Garkavtsev, I.; Jain, R. K., Paracrine regulation of angiogenesis and adipocyte differentiation during in vivo adipogenesis. Circ Res 2003, 93 (9), e88–e97.

48. Sung, H.-K.; Doh, K.-O.; Son, Joe E.; Park, Jin G.; Bae, Y.; Choi, S.; Nelson, Seana Mary L.; Cowling, R.; Nagy, K.; Michael, Iacovos P.; Koh, Gou Y.; Adamson, S. L.; Pawson, T.; Nagy, A., Adipose Vascular Endothelial Growth Factor Regulates Metabolic Homeostasis through Angiogenesis. Cell Metabolism 2013, 17 (1), 61–72.

49. Gharakhanian, R.; Su, S.; Aprahamian, T., Vascular Endothelial Growth Factor-A Deficiency in Perivascular Adipose Tissue Impairs Macrovascular Function. Front Physiol 2019, 10.

50. Landskroner-Eiger, S.; Qian, B.; Muise, E. S.; Nawrocki, A. R.; Berger, J. P.; Fine, E. J.; Koba, W.; Deng, Y.; Pollard, J. W.; Scherer, P. E., Proangiogenic contribution of adiponectin toward mammary tumor growth in vivo. Clinical cancer research : an official journal of the American Association for Cancer Research 2009, 15 (10), 3265–76.

51. Ouchi, N.; Kobayashi, H.; Kihara, S.; Kumada, M.; Sato, K.; Inoue, T.; Funahashi, T.; Walsh, K., Adiponectin Stimulates Angiogenesis by Promoting Cross-talk between AMP-activated Protein Kinase and Akt Signaling in Endothelial Cells*. Journal of Biological Chemistry 2004, 279 (2), 1304–1309.

52. Lee, H. P.; Lin, C. Y.; Shih, J. S.; Fong, Y. C.; Wang, S. W.; Li, T. M.; Tang, C. H., Adiponectin promotes VEGF-A-dependent angiogenesis in human chondrosarcoma through PI3K, Akt, mTOR, and HIF-α pathway. Oncotarget 2015, 6 (34), 36746–61.

53. Huang, C. C.; Law, Y. Y.; Liu, S. C.; Hu, S. L.; Lin, J. A.; Chen, C. J.; Wang, S. W.; Tang, C. H., Adiponectin Promotes VEGF Expression in Rheumatoid Arthritis Synovial Fibroblasts and Induces Endothelial Progenitor Cell Angiogenesis by Inhibiting miR-106a-5p. Cells 2021, 10 (10).

54. Bråkenhielm, E.; Veitonmäki, N.; Cao, R.; Kihara, S.; Matsuzawa, Y.; Zhivotovsky, B.; Funahashi, T.; Cao, Y., Adiponectin-induced antiangiogenesis and antitumor activity involve caspase-mediated endothelial cell apoptosis. Proceedings of the national academy of sciences 2004, 101 (8), 2476–2481.

55. Mahadev, K.; Wu, X.; Donnelly, S.; Ouedraogo, R.; Eckhart, A. D.; Goldstein, B. J., Adiponectin inhibits vascular endothelial growth factor-induced migration of human coronary artery endothelial cells. Cardiovascular Research 2008, 78 (2), 376–384.

56. Elias, I.; Franckhauser, S.; Ferré, T.; Vilà, L.; Tafuro, S.; Muñoz, S.; Roca, C.; Ramos, D.; Pujol, A.; Riu, E., Adipose tissue overexpression of vascular endothelial growth factor protects against diet-induced obesity and insulin resistance. Diabetes 2012, 61 (7), 1801–1813.

57. Wotton, D.; Lo, R. S.; Lee, S.; Massagué, J., A Smad transcriptional corepressor. Cell 1999, 97 (1), 29–39.

58. Seo, S. R.; Ferrand, N.; Faresse, N.; Prunier, C.; Abécassis, L.; Pessah, M.; Bourgeade, M.-F.; Atfi, A., Nuclear retention of the tumor suppressor cPML by the homeodomain protein TGIF restricts TGF-β signaling. Molecular cell 2006, 23 (4), 547–559.

59. Seo, S. R.; Lallemand, F.; Ferrand, N.; Pessah, M.; L’Hoste, S.; Camonis, J.; Atfi, A., The novel E3 ubiquitin ligase Tiul1 associates with TGIF to target Smad2 for degradation. The EMBO journal 2004, 23 (19), 3780–3792.

60. Horie, T.; Ono, K.; Kinoshita, M.; Nishi, H.; Nagao, K.; Kawamura, T.; Abe, Y.; Wada, H.; Shimatsu, A.; Kita, T.; Hasegawa, K., TG-interacting factor is required for the differentiation of preadipocytes. Journal of lipid research 2008, 49 (6), 1224–1234.

61. Björk, C.; Subramanian, N.; Liu, J.; Acosta, J. R.; Tavira, B.; Eriksson, A. B.; Arner, P.; Laurencikiene, J., An RNAi screening of clinically relevant transcription factors regulating human adipogenesis and adipocyte metabolism. Endocrinology 2021, 162 (7), bqab096.

62. Fried, S. K.; Ricci, M. R.; Russell, C. D.; Laferrère, B., Regulation of Leptin Production in Humans. The Journal of Nutrition 2000, 130 (12), 3127S–3131S.

63. Petrescu, A. D.; Grant, S.; Williams, E.; An, S. Y.; Seth, N.; Shell, M.; Amundsen, T.; Tan, C.; Nadeem, Y.; Tjahja, M.; Weld, L.; Chu, C. S.; Venter, J.; Frampton, G.; McMillin, M.; DeMorrow, S., Leptin Enhances Hepatic Fibrosis and Inflammation in a Mouse Model of Cholestasis. The American Journal of Pathology 2022, 192 (3), 484–502.

64. Vivoli, E.; Di Maira, G.; Marra, F., Liver Fibrosis and Leptin. Current Pathobiology Reports 2016, 4 (2), 69–76.

65. Saxena, N. K.; Ikeda, K.; Rockey, D. C.; Friedman, S. L.; Anania, F. A., Leptin in hepatic fibrosis: evidence for increased collagen production in stellate cells and lean littermates of ob/ob mice. Hepatology 2002, 35 (4), 762–71.

66. Liu, Y.; Li, Y.; Liang, J.; Sun, Z.; Wu, Q.; Liu, Y.; Sun, C., The Mechanism of Leptin on Inhibiting Fibrosis and Promoting Browning of White Fat by Reducing ITGA5 in Mice. Int J Mol Sci 2021, 22 (22).

67. Becerril, S.; Rodríguez, A.; Catalán, V.; Mendez-Gimenez, L.; Ramírez, B.; Sainz, N.; Llorente, M.; Unamuno, X.; Gomez-Ambrosi, J.; Frühbeck, G., Targeted disruption of the iNOS gene improves adipose tissue inflammation and fibrosis in leptin-deficient ob/ob mice: role of tenascin C. International Journal of Obesity 2018, 42 (8), 1458–1470.

68. Han, M. S.; White, A.; Perry, R. J.; Camporez, J.-P.; Hidalgo, J.; Shulman, G. I.; Davis, R. J., Regulation of adipose tissue inflammation by interleukin 6. Proceedings of the National Academy of Sciences 2020, 117 (6), 2751–2760.

69. Rehman, K.; Akash, M. S. H.; Liaqat, A.; Kamal, S.; Qadir, M. I.; Rasul, A., Role of Interleukin-6 in Development of Insulin Resistance and Type 2 Diabetes Mellitus. Crit Rev Eukaryot Gene Expr 2017, 27 (3), 229–236.

70. Jin, X.; Yao, T.; Zhou, Z. e.; Zhu, J.; Zhang, S.; Hu, W.; Shen, C., Advanced glycation end products enhance macrophages polarization into M1 phenotype through activating RAGE/NF-κB pathway. BioMed research international 2015, 2015.

71. Sindhu, S.; Thomas, R.; Shihab, P.; Al-Shawaf, E.; Hasan, A.; Alghanim, M.; Behbehani, K.; Ahmad, R., Changes in the adipose tissue expression of CD86 costimulatory ligand and CD163 scavenger receptor in obesity and type-2 diabetes: Implication for metabolic disease. J Glycomics Lipidomics 2015, 5 (134), 2153–0637.100013.

72. Ramsey, S. A.; Klemm, S. L.; Zak, D. E.; Kennedy, K. A.; Thorsson, V.; Li, B.; Gilchrist, M.; Gold, E. S.; Johnson, C. D.; Litvak, V.; Navarro, G.; Roach, J. C.; Rosenberger, C. M.; Rust, A. G.; Yudkovsky, N.; Aderem, A.; Shmulevich, I., Uncovering a Macrophage Transcriptional Program by Integrating Evidence from Motif Scanning and Expression Dynamics. PLOS Computational Biology 2008, 4 (3), e1000021.

73. Maciejewski, M. L.; Arterburn, D. E.; Van Scoyoc, L.; Smith, V. A.; Yancy, W. S., Jr.; Weidenbacher, H. J.; Livingston, E. H.; Olsen, M. K., Bariatric Surgery and Long-term Durability of Weight Loss. JAMA Surg 2016, 151 (11), 1046–1055.

74. Sjöström, L., Review of the key results from the Swedish Obese Subjects (SOS) trial–a prospective controlled intervention study of bariatric surgery. Journal of internal medicine 2013, 273 (3), 219–234.

75. Dai, C.; Liu, Y., Hepatocyte growth factor antagonizes the profibrotic action of TGF-beta1 in mesangial cells by stabilizing Smad transcriptional corepressor TGIF. Journal of the American Society of Nephrology : JASN 2004, 15 (6), 1402–12.

76. Roh, H. C.; Kumari, M.; Taleb, S.; Tenen, D.; Jacobs, C.; Lyubetskaya, A.; Tsai, L. T. Y.; Rosen, E. D., Adipocytes fail to maintain cellular identity during obesity due to reduced PPARγ activity and elevated TGFβ-SMAD signaling. Molecular Metabolism 2020, 42, 101086.

77. Jones, J. E. C.; Rabhi, N.; Orofino, J.; Gamini, R.; Perissi, V.; Vernochet, C.; Farmer, S. R., The Adipocyte Acquires a Fibroblast-Like Transcriptional Signature in Response to a High Fat Diet. Scientific Reports 2020, 10 (1), 2380.

78. Dalmas, E.; Toubal, A.; Alzaid, F.; Blazek, K.; Eames, H. L.; Lebozec, K.; Pini, M.; Hainault, I.; Montastier, E.; Denis, R. G.; Ancel, P.; Lacombe, A.; Ling, Y.; Allatif, O.; Cruciani-Guglielmacci, C.; André, S.; Viguerie, N.; Poitou, C.; Stich, V.; Torcivia, A.; Foufelle, F.; Luquet, S.; Aron-Wisnewsky, J.; Langin, D.; Clément, K.; Udalova, I. A.; Venteclef, N., Irf5 deficiency in macrophages promotes beneficial adipose tissue expansion and insulin sensitivity during obesity. Nat Med 2015, 21 (6), 610–8.

79. Anvari, G.; Bellas, E., Hypoxia induces stress fiber formation in adipocytes in the early stage of obesity. Scientific Reports 2021, 11 (1), 21473.

80. Crewe, C.; An, Y. A.; Scherer, P. E., The ominous triad of adipose tissue dysfunction: inflammation, fibrosis, and impaired angiogenesis. J Clin Invest 2017, 127 (1), 74–82.

81. Reggio, S.; Rouault, C.; Poitou, C.; Bichet, J.-C.; Prifti, E.; Bouillot, J.-L.; Rizkalla, S.; Lacasa, D.; Tordjman, J.; Clément, K., Increased basement membrane components in adipose tissue during obesity: links with TGFβ and metabolic phenotypes. The Journal of Clinical Endocrinology & Metabolism 2016, 101 (6), 2578–2587.

82. Liu, L. F.; Kodama, K.; Wei, K.; Tolentino, L. L.; Choi, O.; Engleman, E. G.; Butte, A. J.; McLaughlin, T., The receptor CD44 is associated with systemic insulin resistance and proinflammatory macrophages in human adipose tissue. Diabetologia 2015, 58, 1579–1586.

83. Ruggiero, A. D.; Key, C. C.; Kavanagh, K., Adipose Tissue Macrophage Polarization in Healthy and Unhealthy Obesity. Front Nutr 2021, 8, 625331.

84. Mancuso, P.; Bouchard, B., The Impact of Aging on Adipose Function and Adipokine Synthesis. Frontiers in endocrinology 2019, 10, 137–137.

85. Lu, B.; Huang, L.; Cao, J.; Li, L.; Wu, W.; Chen, X.; Ding, C., Adipose tissue macrophages in aging-associated adipose tissue function. The Journal of Physiological Sciences 2021, 71 (1), 38.

86. Tominaga, K.; Suzuki, H. I., TGF-β Signaling in Cellular Senescence and Aging-Related Pathology. Int J Mol Sci 2019, 20 (20).

87. Moore, E.; Allen, J. B.; Mulligan, C. J.; Wayne, E. C., Ancestry of cells must be considered in bioengineering. Nature Reviews Materials 2022, 7 (1), 2–4.

88. Bisogno, L. S.; Yang, J.; Bennett, B. D.; Ward, J. M.; Mackey, L. C.; Annab, L. A.; Bushel, P. R.; Singhal, S.; Schurman, S. H.; Byun, J. S., Ancestry-dependent gene expression correlates with reprogramming to pluripotency and multiple dynamic biological processes. Science advances 2020, 6 (47), eabc3851.

89. Ramamoorthy, A.; Pacanowski, M.; Bull, J.; Zhang, L., Racial/ethnic differences in drug disposition and response: review of recently approved drugs. Clinical Pharmacology & Therapeutics 2015, 97 (3), 263–273.

90. Zhang, Q.; Wang, Y.; Huang, E. S., Changes in racial/ethnic disparities in the prevalence of Type 2 diabetes by obesity level among US adults. Ethnicity & health 2009, 14 (5), 439–57.

91. Gao, H. X.; Regier, E. E.; Close, K. L., Prevalence of and trends in diabetes among adults in the United States, 1988-2012. Journal of Diabetes 2016, 8 (1), 8–9.

92. Hales, C. M.; Carroll, M. D.; Fryar, C. D.; Ogden, C. L., Prevalence of obesity among adults and youth: United States, 2015–2016. 2017.

93. Al-Mulla, F.; Leibovich, S. J.; Francis, I. M.; Bitar, M. S., Impaired TGF-β signaling and a defect in resolution of inflammation contribute to delayed wound healing in a female rat model of type 2 diabetes. Molecular bioSystems 2011, 7 (11), 3006–20.

94. Swigris, J. J.; Olson, A. L.; Huie, T. J.; Fernandez-Perez, E. R.; Solomon, J.; Sprunger, D.; Brown, K. K., Ethnic and racial differences in the presence of idiopathic pulmonary fibrosis at death. Respiratory medicine 2012, 106 (4), 588–93.

95. Layden, J. E.; Cotler, S.; Brown, K. A.; Lucey, M. R.; Te, H. S.; Eswaran, S.; Fimmel, C.; Layden, T. J.; Clark, N. M., Racial differences in fibrosis progression after HCV-related liver transplantation. Transplantation 2012, 94 (2), 178–84.

96. Raygor, V.; Abbasi, F.; Lazzeroni, L. C.; Kim, S.; Ingelsson, E.; Reaven, G. M.; Knowles, J. W., Impact of race/ethnicity on insulin resistance and hypertriglyceridaemia. Diabetes & vascular disease research 2019, 16 (2), 153–159.

97. Mo, Y.-Y. Investigating the role of TGIF in beta cell function and diabetes; University of Mississippi Medical Center JACKSON United States: 2020.

98. Arita, Y.; Kihara, S.; Ouchi, N.; Takahashi, M.; Maeda, K.; Miyagawa, J.; Hotta, K.; Shimomura, T.; Miyaoka, K.; Kuriyama, H., Nishida m, Yamashita S, Okubo K, Matsubara K, Muraguchi M, Ohmoto Y, Matsuzawa Y (1999). Paradoxical decrease of an adipose-specific protein, adiponectin, in obesity. Biochemical and biophysical research communications 257, 79–83.

99. Hotta, K.; Funahashi, T.; Arita, Y.; Takahashi, M.; Matsuda, M.; Okamoto, Y.; Iwahashi, H.; Kuriyama, H.; Ouchi, N.; Maeda, K., Plasma concentrations of a novel, adipose-specific protein, adiponectin, in type 2 diabetic patients. Arteriosclerosis, thrombosis, and vascular biology 2000, 20 (6), 1595–1599.

100. Achari, A. E.; Jain, S. K., Adiponectin, a Therapeutic Target for Obesity, Diabetes, and Endothelial Dysfunction. Int J Mol Sci 2017, 18 (6).

101. Yao, Y.; Xu, X.-H.; Jin, L., Macrophage Polarization in Physiological and Pathological Pregnancy. Frontiers in Immunology 2019, 10.

102. Weisberg, S. P.; McCann, D.; Desai, M.; Rosenbaum, M.; Leibel, R. L.; Ferrante, A. W., Obesity is associated with macrophage accumulation in adipose tissue. The Journal of clinical investigation 2003, 112 (12), 1796–1808.

103. Xu, H.; Barnes, G. T.; Yang, Q.; Tan, G.; Yang, D.; Chou, C. J.; Sole, J.; Nichols, A.; Ross, J. S.; Tartaglia, L. A., Chronic inflammation in fat plays a crucial role in the development of obesity-related insulin resistance. The Journal of clinical investigation 2003, 112 (12), 1821–1830.

104. Lumeng, C. N.; Bodzin, J. L.; Saltiel, A. R., Obesity induces a phenotypic switch in adipose tissue macrophage polarization. The Journal of clinical investigation 2007, 117 (1), 175–184.

105. Lumeng, C. N.; DelProposto, J. B.; Westcott, D. J.; Saltiel, A. R., Phenotypic switching of adipose tissue macrophages with obesity is generated by spatiotemporal differences in macrophage subtypes. Diabetes 2008, 57 (12), 3239–3246.

106. Chylikova, J.; Dvorackova, J.; Tauber, Z.; Kamarad, V., M1/M2 macrophage polarization in human obese adipose tissue. Biomedical papers of the Medical Faculty of the University Palacky, Olomouc, Czechoslovakia 2018, 162 (2), 79–82.

107. Tanaka, M.; Ikeda, K.; Suganami, T.; Komiya, C.; Ochi, K.; Shirakawa, I.; Hamaguchi, M.; Nishimura, S.; Manabe, I.; Matsuda, T.; Kimura, K.; Inoue, H.; Inagaki, Y.; Aoe, S.; Yamasaki, S.; Ogawa, Y., Macrophage-inducible C-type lectin underlies obesity-induced adipose tissue fibrosis. Nat Commun 2014, 5, 4982.

108. Gericke, M.; Weyer, U.; Braune, J.; Bechmann, I.; Eilers, J., A method for long-term live imaging of tissue macrophages in adipose tissue explants. Am J Physiol Endocrinol Metab 2015, 308 (11), E1023–33.

109. Cinti, S.; Mitchell, G.; Barbatelli, G.; Murano, I.; Ceresi, E.; Faloia, E.; Wang, S.; Fortier, M.; Greenberg, A. S.; Obin, M. S., Adipocyte death defines macrophage localization and function in adipose tissue of obese mice and humans. Journal of lipid research 2005, 46 (11), 2347–2355.

110. Nawaz, A.; Tobe, K., M2-like macrophages serve as a niche for adipocyte progenitors in adipose tissue. Journal of diabetes investigation 2019, 10 (6), 1394–1400.

111. Gaens, K. H.; Stehouwer, C. D.; Schalkwijk, C. G., Advanced glycation endproducts and its receptor for advanced glycation endproducts in obesity. Curr Opin Lipidol 2013, 24 (1), 4–11.

112. Ruiz, H. H.; Ramasamy, R.; Schmidt, A. M., Advanced Glycation End Products: Building on the Concept of the “Common Soil” in Metabolic Disease. Endocrinology 2020, 161 (1).

113. Wang, Z.; Wang, D.; Wang, Y., Cigarette Smoking and Adipose Tissue: The Emerging Role in Progression of Atherosclerosis. Mediators Inflamm 2017, 2017, 3102737.

114. Solberg, L. I.; Desai, J. R.; O’Connor, P. J.; Bishop, D. B.; Devlin, H. M., Diabetic patients who smoke: are they different? Ann Fam Med 2004, 2 (1), 26–32.

115. Cai, X.; Chen, Y.; Yang, W.; Gao, X.; Han, X.; Ji, L., The association of smoking and risk of diabetic retinopathy in patients with type 1 and type 2 diabetes: a meta-analysis. Endocrine 2018, 62 (2), 299–306.

116. Sari, M. I.; Sari, N.; Darlan, D. M.; Prasetya, R. J., Cigarette Smoking and Hyperglycaemia in Diabetic Patients. Open Access Maced J Med Sci 2018, 6 (4), 634–637.

117. Choi, S. K.; Kim, C. K.; Jo, D. I.; Lee, M. C.; Kim, J. N.; Choi, H. G.; Shin, D. H.; Kim, S. H., Factors Associated with a Prolonged Length of Hospital Stay in Patients with Diabetic Foot: A Single-Center Retrospective Study. Arch Plast Surg 2017, 44 (6), 539–544.

118. Stenkula, K. G.; Erlanson-Albertsson, C., Adipose cell size: importance in health and disease. Am J Physiol Regul Integr Comp Physiol 2018, 315 (2), R284–R295.

119. Hansson, B.; Moren, B.; Fryklund, C.; Vliex, L.; Wasserstrom, S.; Albinsson, S.; Berger, K.; Stenkula, K. G., Adipose cell size changes are associated with a drastic actin remodeling. Sci Rep 2019, 9 (1), 12941.

120. Kim, K. K.; Sheppard, D.; Chapman, H. A., TGF-β1 Signaling and Tissue Fibrosis. Cold Spring Harb Perspect Biol 2018, 10 (4).

121. Sun, K.; Tordjman, J.; Clement, K.; Scherer, P. E., Fibrosis and adipose tissue dysfunction. Cell Metab 2013, 18 (4), 470–7.

122. Francis, M. P.; Sachs, P. C.; Madurantakam, P. A.; Sell, S. A.; Elmore, L. W.; Bowlin, G. L.; Holt, S. E., Electrospinning adipose tissue-derived extracellular matrix for adipose stem cell culture. J Biomed Mater Res A 2012, 100 (7), 1716–24.

123. Di Caprio, N.; Bellas, E., Collagen Stiffness and Architecture Regulate Fibrotic Gene Expression in Engineered Adipose Tissue. Adv Biosyst 2020, 4 (6), e1900286.

124. Ricard-Blum, S., The collagen family. Cold Spring Harb Perspect Biol 2011, 3 (1), a004978.

125. Karsdal, M. A.; Nielsen, S. H.; Leeming, D. J.; Langholm, L. L.; Nielsen, M. J.; Manon-Jensen, T.; Siebuhr, A.; Gudmann, N. S.; Ronnow, S.; Sand, J. M.; Daniels, S. J.; Mortensen, J. H.; Schuppan, D., The good and the bad collagens of fibrosis - Their role in signaling and organ function. Adv Drug Deliv Rev 2017, 121, 43–56.

126. Johannsen, D. L.; Tchoukalova, Y.; Tam, C. S.; Covington, J. D.; Xie, W.; Schwarz, J.-M.; Bajpeyi, S.; Ravussin, E., Effect of 8 Weeks of Overfeeding on Ectopic Fat Deposition and Insulin Sensitivity: Testing the “Adipose Tissue Expandability” Hypothesis. Diabetes Care 2014, 37 (10), 2789–2797.

127. Strieder-Barboza, C.; Baker, N. A.; Flesher, C. G.; Karmakar, M.; Neeley, C. K.; Polsinelli, D.; Dimick, J. B.; Finks, J. F.; Ghaferi, A. A.; Varban, O. A.; Lumeng, C. N.; O’Rourke, R. W., Advanced glycation end-products regulate extracellular matrix-adipocyte metabolic crosstalk in diabetes. Scientific Reports 2019, 9 (1), 19748.

128. Maeda, N.; Shimomura, I.; Kishida, K.; Nishizawa, H.; Matsuda, M.; Nagaretani, H.; Furuyama, N.; Kondo, H.; Takahashi, M.; Arita, Y.; Komuro, R.; Ouchi, N.; Kihara, S.; Tochino, Y.; Okutomi, K.; Horie, M.; Takeda, S.; Aoyama, T.; Funahashi, T.; Matsuzawa, Y., Diet-induced insulin resistance in mice lacking adiponectin/ACRP30. Nat Med 2002, 8 (7), 731–7.

129. Zeyda, M.; Huber, J.; Prager, G.; Stulnig, T. M., Inflammation correlates with markers of T-cell subsets including regulatory T cells in adipose tissue from obese patients. Obesity (Silver Spring) 2011, 19 (4), 743–8.

130. Buechler, C.; Krautbauer, S.; Eisinger, K., Adipose tissue fibrosis. World journal of diabetes 2015, 6 (4), 548–53.

131. Beltowski, J., Adiponectin and resistin--new hormones of white adipose tissue. Med Sci Monit 2003, 9 (2), RA55–61.

132. Kern, P. A.; Di Gregorio, G. B.; Lu, T.; Rassouli, N.; Ranganathan, G., Adiponectin Expression from Human Adipose Tissue. Diabetes 2003, 52, 1779–1785.

133. Wang, J.; Leclercq, I.; Brymora, J. M.; Xu, N.; Ramezani-Moghadam, M.; London, R. M.; Brigstock, D.; George, J., Kupffer cells mediate leptin-induced liver fibrosis. Gastroenterology 2009, 137 (2), 713–23.

